# Morphological innovation without gene co-option: the *Drosophila* sex comb evolved via changes in developmental tempo and energy metabolism

**DOI:** 10.64898/2026.02.13.705815

**Authors:** Ben R. Hopkins, Olga Barmina, Xinying Wang, Mandy M. Situ, Haley A. Bolanos, Shizhan Nie, Artyom Kopp

## Abstract

The individualization of serially repeated homologs is one route through which novel traits are thought to evolve. Under this model, a repeated character—like a limb, digit, or sensory bristle—is individualized from its homologs by changes in the regulatory apparatus (‘character identity network’, ChIN) that specifies its development. Individualization then enables downstream gene networks that build the repeated character to diverge from one another in different parts of the body, ultimately allowing new phenotypic endpoints to be reached. Despite this model’s intuitive appeal, the genetic mechanisms through which new ChINs rewire trait-building gene networks remain largely uncharacterized. A promising system in which to study this process is the *Drosophila* sex comb. Found in a sublineage of *Drosophila* species, the sex comb is a recently evolved, male-specific innovation that evolved from a more evolutionarily ancient precursor—the mechanosensory (MS) bristle—following the gain of a novel ChIN centered on the sex determination gene *dsx* and HOX gene *Scr*. Here, we use time-series single-cell RNA-seq to show that rather than co-opting new genes, this new ChIN orchestrates quantitative and heterochronic changes in the ancestral MS bristle transcriptome. These changes affect gene modules that control energy metabolism, endoreplication, and actin dynamics. The net effect of these changes is an organ-specific shift in developmental rate, leading to accelerated growth in sex comb teeth. Collectively, our work suggests that morphological innovation can proceed without the co-option of new genes into downstream trait-building networks and instead through metabolically driven differences in developmental rate between serial homologs.

## Introduction

Whether it’s the fur of mammals, the tetrapod limb, or the firefly’s light organ, ‘innovations’ have evolved repeatedly across the tree of life, unlocking new ecological opportunities and evolutionary trajectories for the lineages in which they arise. Despite the apparent ubiquity of innovations, understanding how the evolutionary process can generate entirely new body parts and biological functions remains one of biology’s major challenges (1–6). Previous work has emphasized the individualization of serial homologs as a pathway through which novel traits evolve. In this scenario, the development of a repeated character—such as a wing, limb, or digit—is underpinned by a three-tier genetic logic, where the middle tier is occupied by a ‘character identity network (ChIN)’ that takes in upstream positional information and relays it into the downstream activation of networks of ‘realizer’ genes that build a particular character (1). A prediction of this model is that the modification of a ChIN in one part of the body leads to a regulatory separation between homologous characters in other parts of the body. This regulatory separation in turn allows the realizer networks downstream of the modified and ancestral ChINs to evolve independently from one another, ultimately giving rise to qualitatively different phenotypic endpoints—for example, one wing pair remaining as a wing pair while the other is transformed into a beetle’s elytra or a fly’s haltere (7). This is an attractive model for the origin of new traits, but it would be strengthened by case studies that identify how the evolutionary modification of ChINs rewires the downstream gene networks they control.

In principle, there are three broad modes of gene expression change that could occur in an evolving realizer network. First, there could be qualitative changes in expression, where genes that aren’t expressed in the ancestral homolog are recruited from other tissues. At its most extreme, this recruitment may extend beyond single genes to the co-option of entire gene modules active in other cell types. Intuitively, module co-option provides a compelling mechanism through which large-scale phenotypic shifts can be rapidly achieved: complex networks of interacting genes assembled in one tissue over long expanses of evolutionary time can be redeployed wholesale through simple changes in the spatial activity of their upstream regulators. Indeed, there is now a bank of elegant examples that implicate module co-option in the gain of novel traits (8–10). But studies have also shown that non-qualitative changes in gene expression—changes in either the level (‘quantitative’) or the timing (‘heterochronic’) of expression of existing genes—can also be responsible for major anatomical differences between closely related species. In mammals, for example, heterochronic change in the expression of the transcription factor SATB2 has been linked to the origin of the corpus callosum, an axon tract that connects the left and right cerebral cortices in eutherian mammals (11, 12). While both qualitative and non-qualitative changes in gene expression offer potential routes to innovation, the relative importance of each remains unclear.

With its recent evolutionary origin and presence in a genetically tractable system, the *Drosophila* sex comb has emerged as a model system for studying the genetic basis of evolutionary innovation (5). Absent in most *Drosophila* species and restricted to members of the *Sophophora-Lordiphosa* radiation, the sex comb is a structure on the male foreleg tarsus that promotes male copulatory success through grasping or stimulating females during mating (13–17). Composed of structures called ‘teeth’, the sex comb is a modified serial homolog of one of the many transverse rows of mechanosensory bristles (TBRs) that cover the forelegs of *D. melanogaster* and most other Diptera species (**Fig. 1A**)(18). Those modifications are threefold. First, the morphology of the bristles is changed to become the much thicker, more elongated, blunter, and more heavily melanized sex comb teeth. Second, the number of teeth is expanded relative to the homologous TBR in females. Third, the whole row is rotated 90° such that it sits in a longitudinal, rather than transverse, orientation—a process driven by a morphogenetic rearrangement of the surrounding epithelial cells (18).

**Figure 1.**
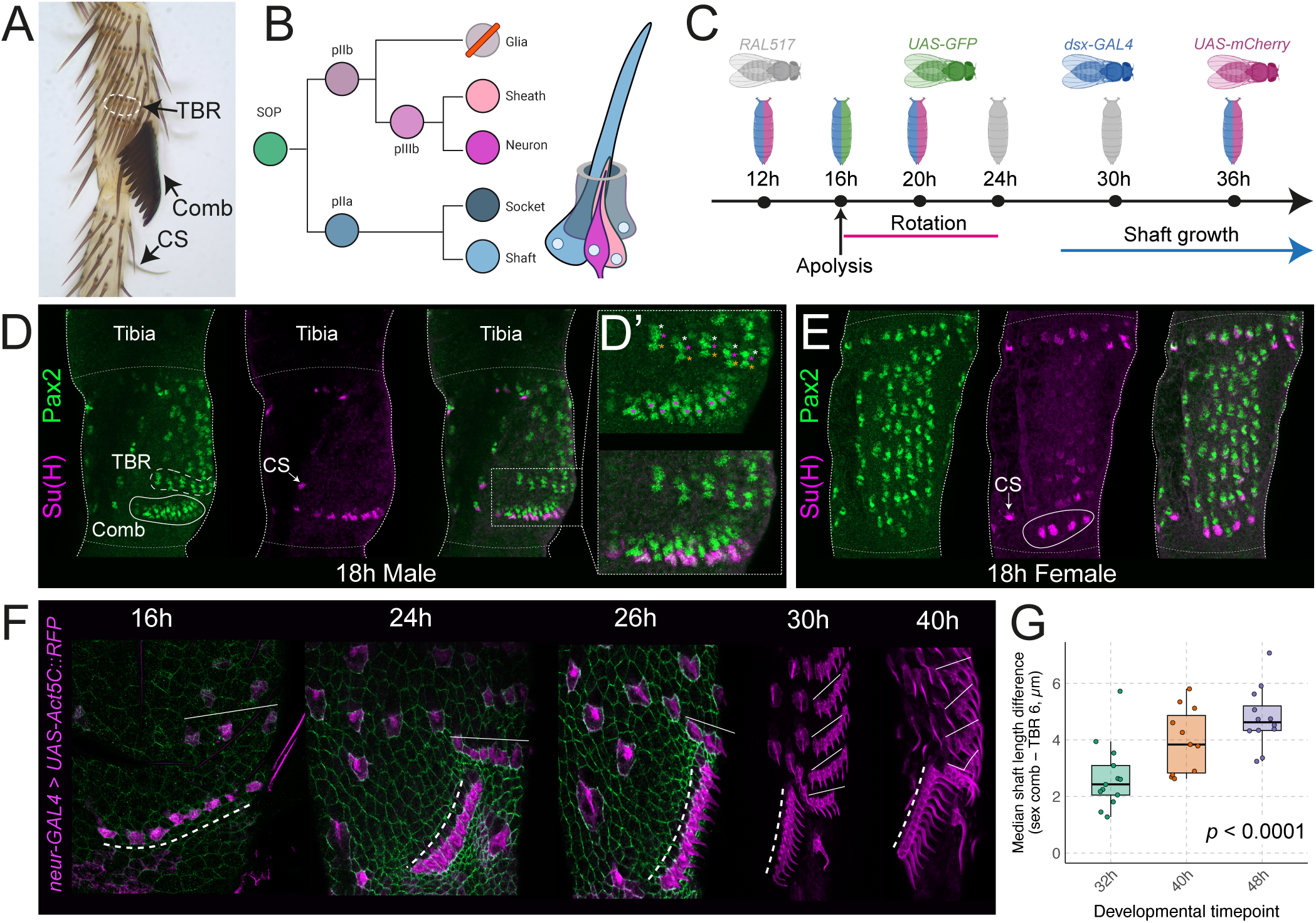
The bristle-to-tooth transformation is achieved through accelerated growth. (A) The first tarsal segment (ta1) of the male foreleg with the sex comb (‘comb’), a chemosensory bristle (‘CS’), and a transverse bristle row (‘TBR’) highlighted. TBRs are composed of mechanosensory (MS) bristles. (B) A diagram of the cell lineage that gives rise to the four cell types that make a bristle. CS bristles develop according to this same general blueprint but in this case multiple neurons are generated. The glia cell undergoes apoptosis (34). Colors map onto those used in Fig. 2. SOP = sensory organ precursor. (C) A time-course of sex comb development. Labelled timepoints correspond to the 6 timepoints sampled for single-cell RNA-sequencing. Colors denote the genotypes sequenced at each stage as part of the single-cell experiments outlined later in the paper. Times are given in hours after puparium formation (APF). (D) Stainings of an 18h APF male ta1 with antibodies against Pax2 and Su(H) (see Supp. Fig. 1 for an image that prioritizes the view of the TBRs rather than the comb). The tibia/ta1 and ta1/ta2 boundaries are marked by horizontal dotted white lines. Su(H) is spe-cifically expressed in socket cells and at this timepoint is largely restricted to the comb teeth and CS bristles. An example of the latter is labelled as ‘CS’. In each developing bristle, the shaft cell can be identified both by its position, falling proximal to the most distal cell (the socket), and by its elevated accumulation of Pax2 (31). Pax2 is ultimately restricted to the sheath and shaft cells in the post-mitotic, differentiative phase of bristle development, but is expressed in all cells of the bristle lineage in the mitotic stage when the cell fates are still being specified (31). In the top image of (D’), three nuclei can be seen stained by anti-Pax2 in each bristle in the most distal TBR (shown with asterisks: orange=socket, magenta=shaft, white=sheath/neuron). In the sex comb, shaft cells, which have elevated levels of Pax2, are marked with magenta asterisks. At least three nuclei are present in each tooth. Whether the non-shaft/non-socket Pax2+ cell is the undivided pIIb cell or a sheath cell with the neuron having lost Pax2 expression is unclear from this staining. The bottom image is the same as the top but also showing the expression of Su(H). (E) As (D) but in a female. The TBR homologous with the sex comb is circled. (F) Shaft growth visualized by an RFP-tagged Actin5C expressed under the control of the bristle driver *neur*-GAL4. Dashed lines show the position of the sex comb and straight lines show TBRs. The green staining in earlier timepoints is anti-Flamingo, which marks cell boundaries. (G) The difference in length between sex comb and MS shafts (comb – MS) at different developmental timepoints. At 26h APF, bristles/ teeth were too short to measure. Each dot represents the difference in the median length of bristles in TBR 6 and the sex comb in a single leg. The p-value is based on a likelihood ratio test comparing linear mixed effects models with and without the interaction term between Organ (comb/TBR 6) and Age (see Supp. Fig. 2). The significant positive interaction indicates that the rate of shaft growth is accelerated in the sex comb.

The sex comb teeth and TBR mechanosensory (MS) bristles are themselves serially homologous to a range of other sensory organs present in the fly, including chemosensory (CS) taste bristles, campaniform sensilla, and chordotonal organs (18–21). Each of these multicellular organ classes develops according to the same general program in which a sensory organ precursor (SOP) cell undergoes multiple rounds of asymmetric division to generate a neuron and accessory cells (**Fig. 1B**). The specific differentiation trajectories that SOP-descendent cells follow have been repeatedly modified throughout evolutionary history, giving rise to new accessory cell types that confer new sensory capabilities, as well as new non-sensory structures like butterfly scales (19, 22). In the case of sensory bristles and sex comb teeth, the critical innovation was the origin of the shaft cell (the ‘trichogen’). Born through the same division as the socket cell that anchors it, the shaft cell assembles bundles of polymerizing actin filaments that are attached to and push out against the plasma membrane to form a hair-like projection (reviewed in 23). A thickened cuticle eventually forms around the projection, the actin bundles break down, and the shaft cell undergoes cell death (24–27). While the developmental genetic changes responsible remain uncharacterized, the shaft cell itself appears to have been repeatedly modified between bristle classes (*e.g.,* mechanosensory vs chemosensory bristles vs sex comb teeth) leading to striking differences in morphology.

The ChIN that individualizes the sex comb from the surrounding TBRs consists of two transcription factors: the HOX gene *Sex combs reduced* (*Scr*) and the principal effector of sexual differentiation, *doublesex* (*dsx*)(28, 29). Through a molecular cascade that starts with a sex chromosome-counting system, *dsx* undergoes sex-specific alternative splicing to generate male- and female-specific isoforms that share a common DNA-binding domain but differ in the sequence of their N-terminus (reviewed in 30). This enables Dsx to exert different regulatory effects on the same target genes in each sex. The expression of *dsx* in the sex comb-bearing region of the first tarsal segment is a derived condition and its gain coincides with the evolution of the sex comb (28). The activation of *dsx* expression is under the control of *Scr*, a gene that is ancestrally expressed at low, sexually monomorphic, and consistent levels throughout the tarsus (28, 29). However, in sex comb-bearing species, a novel autoregulatory feedback loop between *Scr* and the male-specific isoform of *dsx* has evolved, which serves to elevate the expression of both transcription factors in the male sex comb-bearing region (28). It’s this novel feedback loop that defines the individuated sex comb ChIN, providing sex comb teeth with a regulatory apparatus that is distinct from the apparatus active in the surrounding MS bristles.

Here, we focus our attention on changes in the SOP lineage that underlie the bristle-to-tooth transformation. We use time-series single-cell RNA-sequencing (scRNA-seq) to show that the *dsx-Scr* ChIN transforms MS bristles into teeth not through the co-option of genes from other organs or cell types but instead via heterochronic and quantitative changes in gene expression in the MS bristle shaft morphogenetic program. These changes include a strong and early upregulation of genes involved in energy metabolism, protein synthesis, and certain actin-related processes. We also show that the cells of the sex comb undergo additional, male-specific endocycles to reach higher ploidy levels. These changes shift the tempo of shaft development, leading to an acceleration of bristle growth in sex comb teeth relative to their MS homologs. Collectively, our work shows that novel ChINs needn’t co-opt new genes to build novel traits but instead can tune existing cellular processes to reach new phenotypic endpoints.

## Results

### Earlier specification is not sufficient for the bristle-to-tooth transformation

To understand how the morphological differences between sex comb teeth and MS bristles arise, we began by assembling a time course of their development (**Fig. 1C**). Prior to ∼16h APF the pupal cuticle remains attached to the epidermis, which precludes antibody staining. At 18h APF, soon after the separation of cuticle and epidermis (‘apolysis’), antibody stainings against Pax2 showed that the SOP-descendant cells have undergone the final divisions that give rise to the socket and shaft cells in both the sex comb and neighbouring TBRs (**Fig. 1D**). There was, however, a clear difference in the progress of differentiation between these organ classes. At ∼18h APF, commitment to the socket lineage—inferred from the presence of the socket-defining transcription factor Su(H)—was clearest in the sex comb and a handful of other organs, the positions of which were consistent with those of CS organs and possibly campaniform sensilla. In contrast, in most TBRs Su(H) was largely undetectable at this timepoint (**Fig. 1D**), although weak Su(H) expression was detectable in the most proximal TBR (**Supp. Fig. 1**). Given that the shaft derives from the same division as the socket, it is likely that the shaft cell also begins to differentiate earlier in sex comb teeth than in the TBRs. Consistent with this, at 18h APF we observed that Pax2 accumulation in the shaft cells was high in the comb relative to the most distal TBR (**Fig. 1D’**), a sign that comb shaft cells are further along in their differentiation than their MS homologs (31).

Earlier specification of sex comb shaft and socket cells could provide a simple mechanism for generating many of the phenotypic differences that distinguish sex comb teeth from MS bristles because a longer developmental window offers more time in which to grow. To test this, we ran the same stainings on female first tarsal segments at 18h APF. Just as in males, we found that Su(H) staining was substantially stronger in the cells of the most proximal and most distal TBR (**Fig. 1E**). Unlike in males, however, the most distal TBR in females—the sex comb homolog—is morphologically indistinguishable from the other TBRs in the first tarsal segment (e.g., 18). We therefore conclude that earlier specification is not sufficient for the bristle-to-tooth transformation and is instead a positional effect, with bristle cells proximally and distally closest to joints differentiating first.

### Accelerated growth drives the bristle-to-tooth transformation

If early specification is not sufficient to achieve the elongated, thickened morphology of a sex comb tooth, we reasoned that perhaps the rate of shaft growth is accelerated in the sex comb relative to the surrounding MS bristles. To test this, we expressed an RFP-tagged form of the actin Act5C under the control of the bristle driver *neur-*GAL4 and measured the length of the sex comb teeth and the MS bristles in the adjacent TBR at different stages during pupal development. In both MS bristles and sex comb teeth, shaft growth was first detectable at 26h APF, soon after the completion of rotation (**Fig. 1F**). Measuring at 32h, 40h, and 48h APF, we found that the difference in shaft length between sex comb teeth and MS bristles increased through time (linear mixed effects model, LRT, χ^2^(2) = 51.609, *p*<0.0001; **Fig. 1G**; see **Supp. Fig. 2** for model estimates). This temporal increase in the size difference between the two organ classes indicates a shift in developmental tempo, with the rate of shaft growth accelerated in the sex comb relative to MS bristles.

### Transcriptomic differences between sex comb and mechanosensory bristle cells appear early in differentiation

In search of genetic determinants of the divergent growth rates between sex comb teeth and MS bristles, we assembled a time-series scRNA-seq dataset of the sensory organ populations present in the developing first tarsal segment. Sequencing this segment at 12h, 16h, 20h, 24h, 30h and 36h APF, our dataset spans the full set of divisions in the SOP lineage and much of the growth of the shafts (**Fig. 2**; see lineage schematic in **Fig. 1B**) but stops at the point when the epicuticle starts to form (36h APF, 32). We were unable to dissociate the sex comb-bearing region after this timepoint. Consequently, our dataset does not capture the activation of the melanisation pathway. SOP-derived cells form contiguous clusters in the early datasets (12h-20h) with the intermediary progenitor cell populations of pIIa, pIIb, and pIIIb present and identifiable based on the expression of *cato, ham, scrt, spdo, ttk, insv, ase,* and *sens* (see **Supp. Figs. 3-8** for markers used in the annotation of each dataset). At 12h and 16h, we also recover the *repo^+^*glia cell population born from the division of pIIb. Previous work in the notum has shown that this glial cell undergoes apoptosis mediated by the proapoptotic gene *grim* (33, 34). Consistent with an equivalent apoptotic fate, we find that our pIIb-derived glia population is enriched for *grim* and that these cells are absent from our datasets from 20h onwards. Differences between the socket and shaft cells of the sex comb and MS bristles were apparent from the earliest stage we looked at (12h APF) and persisted through to the latest timepoint (36h APF) (**Fig. 2**). The same is true for the CS bristle cells. Given that we detected no discernible shaft growth prior to 26h APF, this suggests that the transcriptomic distinctiveness of the comb (and CS) cells long predates their morphological distinctiveness.

**Figure 2.**
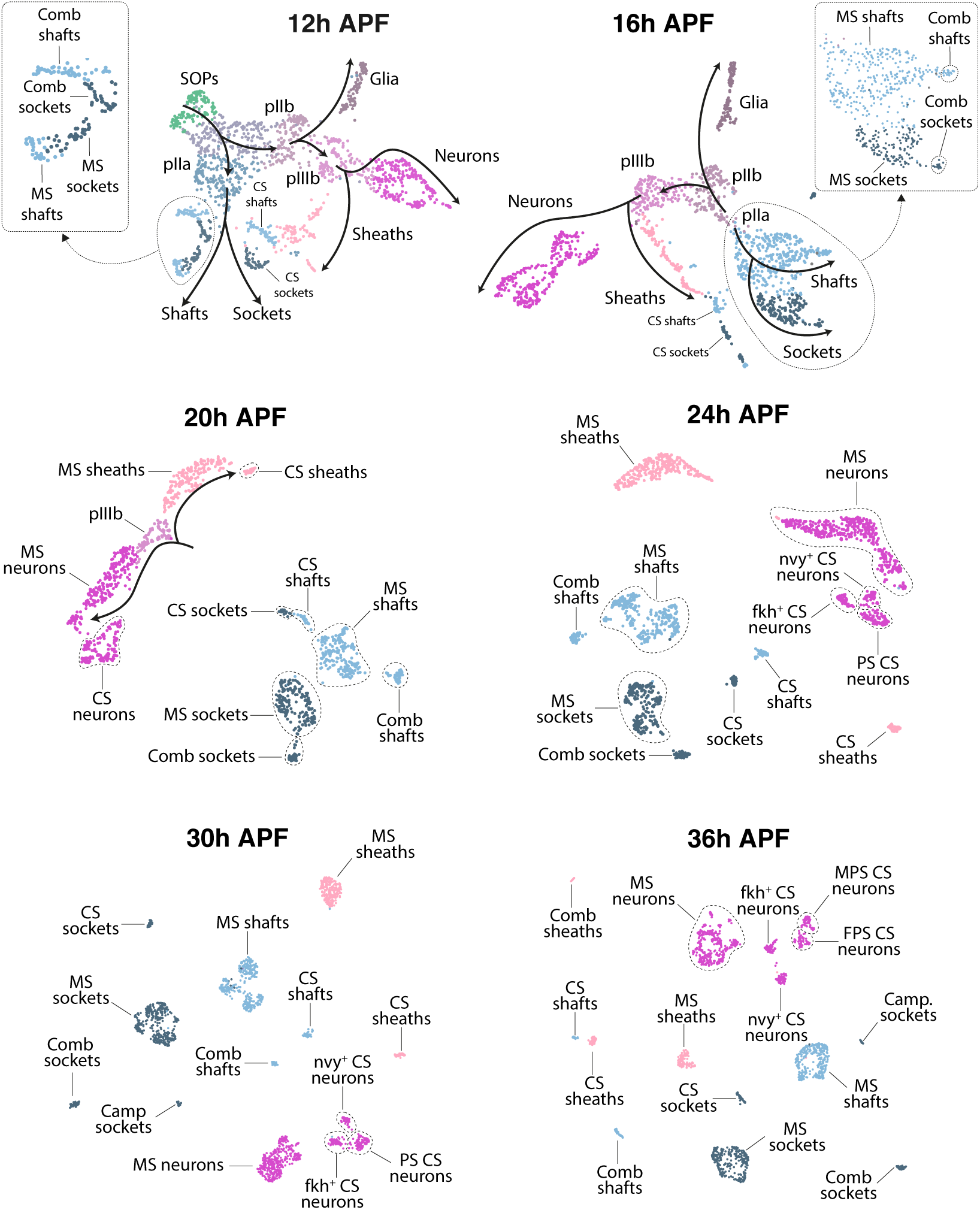
Sex comb cells are transcriptomically distinct before they become morphologically distinct. UMAP plots of the SOP-descendant cells in each of the 12h, 16h, 20h, 24h, 30h, and 36h after puparium for-mation (APF) datasets. Each UMAP was built using the same set of clustering parameters. Progenitor cells (SOPs, pIIa, pIIb, and pIIIb) can be detected in some of the datasets prior to 24h. At all timepoints, comb and chemosensory (CS) socket and shaft cells can be identified. Cells are coloured according to the cell type they belong to and a consistent color scheme, the same used in the Fig. 1B schematic, is used across timepoints. Genes used to guide the annotation are given in Supp. Figs. 3-8. Mechanosensory (MS) bristles are ultimately considerably more numerous than sex comb teeth in the first tarsal segment. The underrepresentation of MS shafts and sockets relative to their sex comb homologues at 12h and 16h APF is because some of the MS pIIa cells have yet to divide. Of those that have, some of the descendant sockets and shafts are less differentiated than in the sex comb and therefore fail to cluster separately from one another. Note that we failed to identify a distinct comb sheath population. Sex comb sheaths are likely insufficiently distinct from their MS homologues to cluster separately. The MS neuron clusters contain MS neurons that innervate MS bristles, chemosensory bristles, campaniform sensilla, and the sex comb. These subpopulations can be separately identified within the MS neuron clusters—with increasing ease at later timepoints—based on the expression of marker genes reported by Hopkins *et al.* (81). While expressing fluorescent reporters under the control of *dsx*-GAL4 aided our identification of sex comb cells in some of our datasets (Fig. 1C), fluorescent reporter expression was not the basis of their distinct clustering. Sex comb cells clustered separately even when *GAL4* and *mCherry* transcripts were removed (Supp. Fig. 9). MS = Mechanosensory, CS = Chemosensory, Camp. = Campaniform sensillum, PS = Pheromone-sensing, MPS = Male pheromone-sensing, FPS = Female pheromone-sensing. Neuron nam-ing convention follows that used by Hopkins *et al.* (81).

### Transcriptomic divergence between mechanosensory and sex comb cells increases with developmental time

We next sought to understand how the transcriptomes of sex comb and MS cells diverge from one another during development. To explore this, we sequenced the first tarsal segment in females at two of the timepoints we sequenced in males: 12h and 24h APF. At 12h APF, cells annotated as sex comb shafts and sockets in the male dataset co-clustered with female cells in an integrated male-female dataset (**Fig. 3A**). At 24h APF, however, male-derived sex comb cells clustered separately from female cells (**Fig. 3B**). This difference in sex comb cell properties between the two datasets suggests that the signal of transcriptomic distinctiveness in sex comb cells is initially driven by features that are shared with a subset of female cells, such as positional identity, but later by features that are male- and comb-specific.

**Figure 3.**
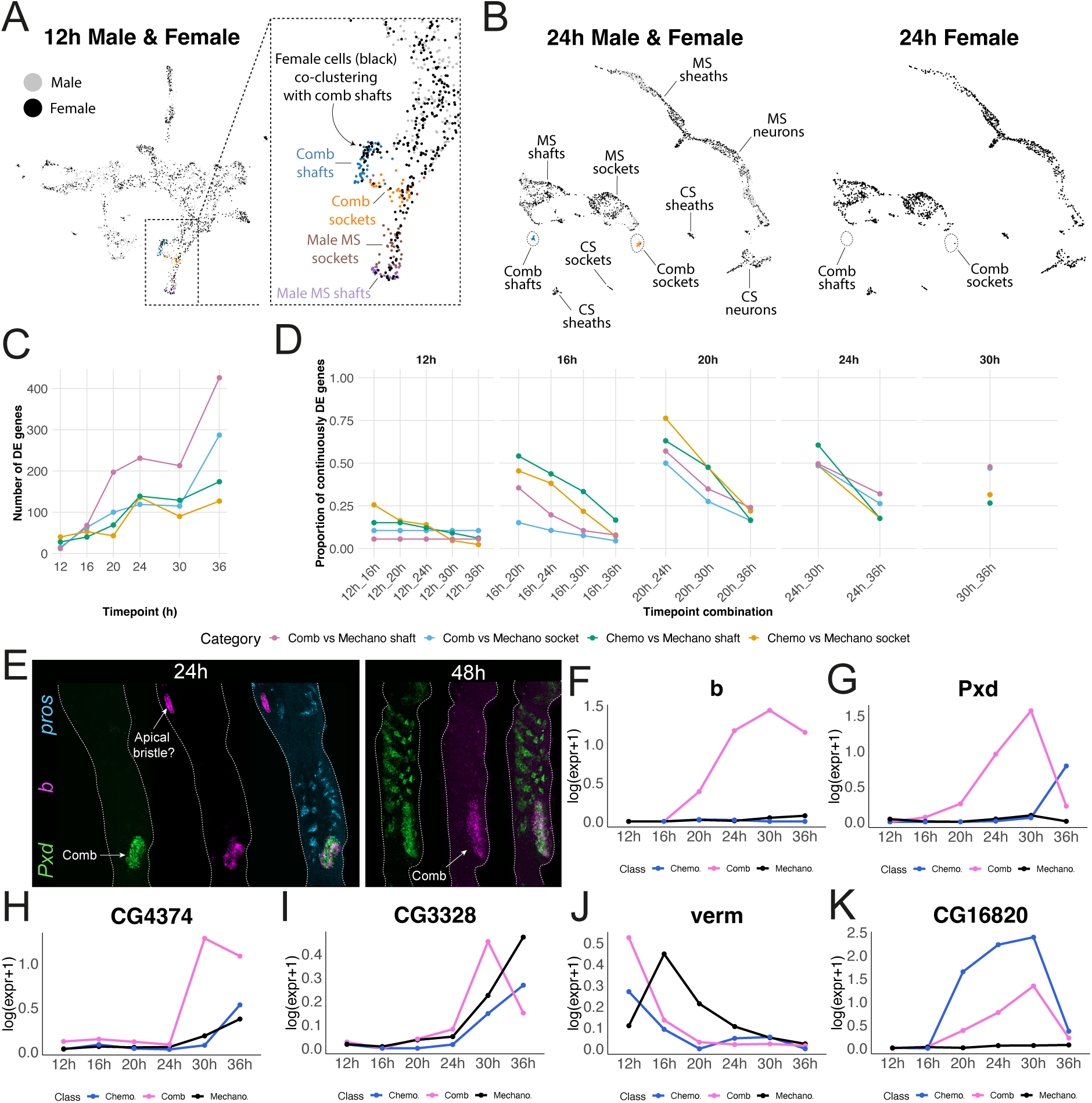
Sex comb cells exhibit quantitative and heterochronic shifts in the expression of conserved bristle genes. (A) A UMAP of an integrated dataset of 12h APF male and female sensory organ lineage cells showing that female cells co-cluster with male sex comb shaft and socket cells. Female cells are coloured black and male cells grey, except for male mechanosensory (MS) shafts (purple), male MS sockets (brown), comb shafts (blue), and comb sockets (orange). For male cells, the annotation for each cell is transferred over from the annotations made on the unintegrated, stand-alone male dataset (as shown in Supp. Fig. 3). (B) UMAPs of an integrated dataset of 24h APF male and female sensory organ lineage cells showing cells from both sexes combined (left) and the female cells alone (right). Cells were annotated in the individual, sex-specific datasets before integration (shown in Supp. Fig. 6 for the male). The position of the sex comb shaft and socket clusters is shown in both datasets. (C) The number of differentially expressed genes (DEGs) between sex comb sock-ets and MS sockets, sex comb shafts and MS shafts, chemosensory (CS) sockets and MS sockets, and CS shafts and MS shafts at each sampled time point. (D) The proportion of DEGs that continue to be differentially expressed at each subsequent timepoint. Values on the x-axis give the start and end dataset being considered (e.g., 16h-30h requires a gene to be differentially expressed at 16h, 20h, 24h, and 30h). (E) HCR in situ stain-ing of 24h APF and 48h APF legs with probes against *Pxd*, *b*, and *pros*. *pros* is used here as a bristle marker, being expressed in the sheath cell of every bristle (and the neurons of CS bristles)(81). The position of the sex comb is labeled, as is the likely position of the pre-apical bristle in the distal tibia. Although *b* shows a qualita-tive difference in expression between sex comb and MS cells, we failed to detect any effect of *b* on sex comb development using a combination of hypomorphic GAL4 lines, deficiency lines, and a molecular null of *b* (Supp. Fig. 11B). Moreover, a sex comb-like darkening of the bristle is the opposite of what we would expect to see on activation of *b* expression—it’s the loss of *b* function that’s associated with darkening of the cuticle (82). (F-K) Time-series expression plots for genes showing qualitative (*b*) and heterochronic (*Pxd, CG4374, CG3328, verm*, *CG16820*) expression differences between sex comb, CS bristle, and MS bristle shaft cells.

We recovered a similar signal of increasing transcriptomic distinctiveness when performing a differential expression analysis between sex comb cells (shafts and sockets, separately) and their homologous cell types in MS bristles at each of the 6 timepoints in males. As a reference, we performed the same analysis between MS cells and CS cells, which, like sex comb cells, also start differentiating earlier (e.g., **Fig. 1D**). We observed that the number of differentially expressed genes (DEGs) increased with time for both the sex comb vs MS and CS vs MS cell comparisons, suggesting that both organs show increasing divergence from their MS homologs over the course of differentiation (**Fig. 3C**). However, from 20h APF onwards, the degree of divergence from the MS homolog was markedly higher for sex comb shafts than it was for CS shafts—we recovered many more DEGs in the comb vs MS shaft comparisons than we did in the CS vs MS comparison (Fig. 3C). A relatively small share of genes that were differentially expressed at an earlier timepoint continued to be differentially expressed at later timepoints (**Fig. 3D**). This suggests that few differences persist over the entire time course of development. Indeed, of the 784 DEGs we identified between sex comb shafts and MS bristle shafts across all timepoints, *dsx* was the only gene differentially expressed in each. Only five genes, including the actin-binding protein *flr* (see below), were differentially expressed at five of the six timepoints.

### The transcriptomic differences between sex comb cells and mechanosensory cells are quantitative and heterochronic rather than qualitative

We next sought to test for genes that show qualitative differences in expression between MS cells and sex comb cells. As a control, we began by looking at CS bristles. As an organ class with a much deeper evolutionary history, we reasoned that the potential for qualitative differences was highest here. Searching through the CS vs MS DEGs, we found several credible instances of qualitative differences and validated their CS specificity using HCR *in situ*. These genes showed persistent expression across timepoints in CS cells with essentially no expression in MS cells (*Ance-3* in sockets; *mtg, CG32040,* and *CG14280* in shafts; *NLaz* in sheaths; **Supp. Fig. 10**). By contrast, of the 784 DEGs between sex comb shafts and MS bristle shafts, we found only a single convincing candidate for qualitative gain: the aspartate decarboxylase gene *black* (*b*). HCR *in situ* stainings confirmed that *b* was restricted to the sex comb in the male first tarsal segment across all the timepoints we sampled (24h, 30h, and 48h) and absent from the homologous region in females (**Fig. 3E,F; Supp. Fig. 11A**). However, we failed to detect any effect of *b* on sex comb development using multiple reagents, including molecular nulls of *b* (*b[1]/Df*; **Supp. Fig. 11B**), and therefore failed to find any evidence of functionally significant qualitative differences in expression between the ancestral and derived cell types.

If the changes in gene expression between comb and MS cells are not qualitative then they must affect genes that are shared between the two organ classes, reflecting changes in either the level or timing of expression. Indeed, we found widespread evidence of both. In some cases, genes exhibiting heterochronic changes initially appeared to be specifically expressed in sex comb cells but then lost their comb-specificity at later timepoints (*e.g., Pxd*), while many of the differences between comb and MS cells were transient and appeared at different timepoints (**Fig. 3E-K**). Moreover, not all DEGs exhibit heterochrony. In other cases, we detected quantitative changes in expression, with the temporal trajectories being similar between cell types across organ classes but with expression generally higher in the comb, as in the case of many ATP synthase subunits (**Supp. Fig. 12**). Collectively, therefore, the patterns of expression change on display were not as simple as if the whole sex comb differentiation program runs ahead of MS cells in the earlier differentiating organ classes (CS bristles and comb), instead pointing to relatively fine-scale, gene-specific modulation of expression level and timing.

### Changes in metabolic gene expression are associated with increased growth in the sex comb

To understand which genes are undergoing quantitative and heterochronic changes in expression in sex comb cells, we used DPGP (Dirichlet Process Gaussian Process mixture model, 35) to cluster differentially expressed genes. This method uses a Gaussian process to model the temporal trajectories of gene expression across time and a Dirichlet process to cluster those trajectories. Using this approach, we recovered 14 clusters in shafts that collectively describe multiple waves of gene expression changes through time. In 10 of these clusters, we identified significant enrichment of at least one biological process term in a GO analysis **(Fig. 4A; Supp. Fig. 13**). These terms largely centered on the processing and production of proteins and ATP. On the protein side, an early upregulation (12h and 16h) of ribosome biogenesis genes (GO:0042254; **Fig. 4B**) in sex comb shafts was followed by later peaks in protein localization to the endoplasmic reticulum (GO:0070972; 20h-30h; **Fig. 4C**), protein maturation (GO:0051604; 30h), and glycosylation (GO:0070085; 36h). On the energy side, we saw persistent upregulation in sex comb shafts of terms including ATP metabolic process (GO:0046034), oxidative phosphorylation (GO:0006119; **Fig. 4D**), generation of precursor metabolites and energy (GO:0006091), and aerobic respiration (GO:0009060). The enrichment of these energy- related terms amounted to a near total upregulation of the entire electron transport chain machinery, including ATP synthase subunits, cytochrome c oxidase (COX) subunits, and NADH dehydrogenases. Consistent with a boost to the energy production machinery, we also detected strong upregulation of nuclear-encoded mitochondrial genes (GO:0005739; **Fig. 4E**), which may reflect an increase in the size of the mitochondrial compartment in the sex comb cells. Upregulation of all these gene modules in the sex comb was more pronounced in the shaft cells than the sockets, mirroring the cell types in which the gross morphological differences between sex comb teeth and MS bristles are most pronounced.

**Figure 4.**
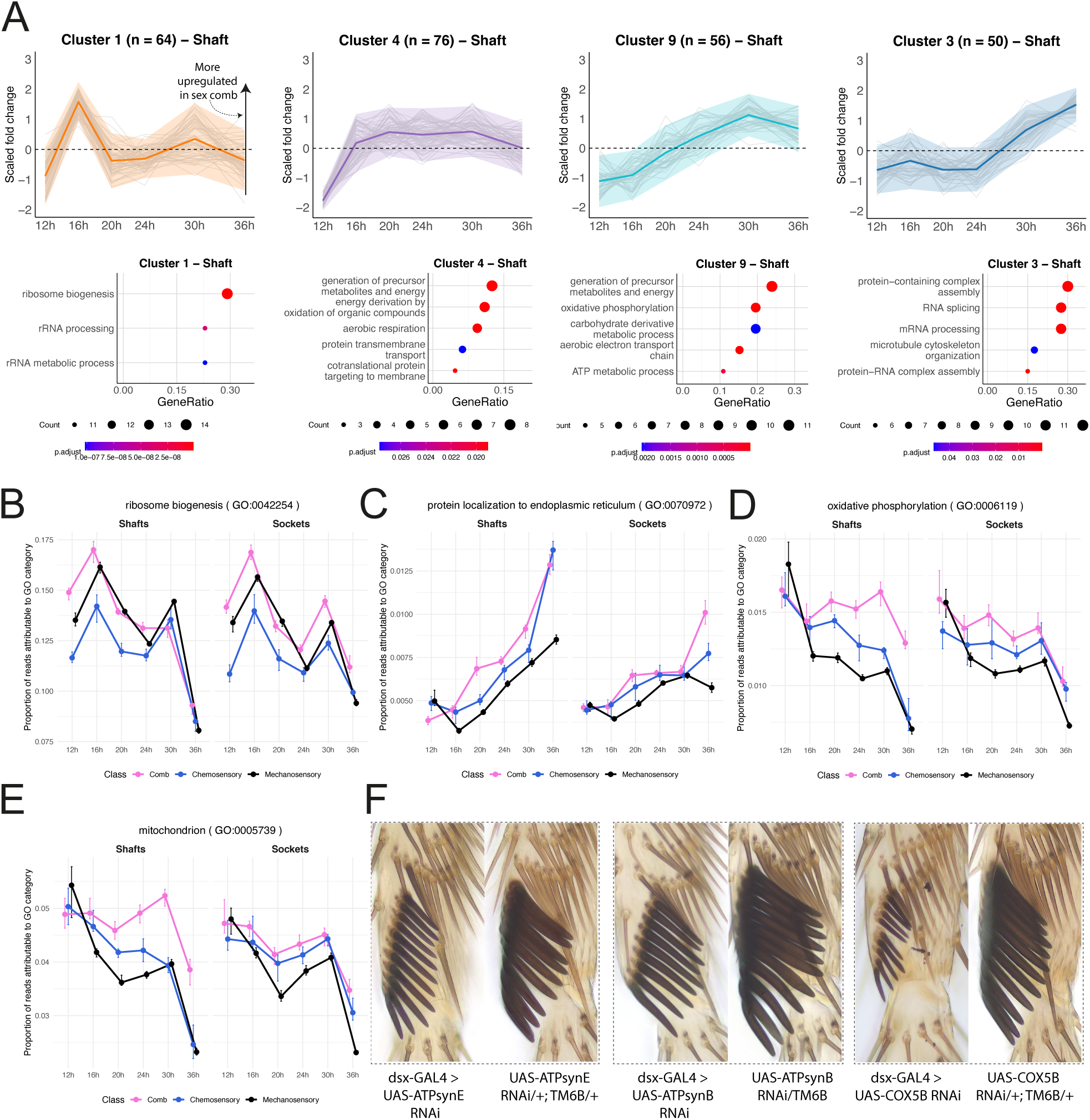
Quantitative and heterochronic augmentation of metabolic gene modules in sex comb shafts. (A) Expression trajectories for genes found to be differentially expressed between sex comb shafts and their mechanosensory (MS) homologs. Using DPGP (35), differentially expressed genes are clustered based on how the magnitude of the comb-MS difference in expression changes across developmental timepoints. 4 of 14 clusters are shown, with the remainder in Supp. Fig. 13. Each expression plot shows the trajectory of each gene in the cluster (gray lines) as well as the cluster mean (thick, colored line). The ribbon denotes ± 2 S.D. The number of genes in the cluster is given in the plot title. The values plotted are the log2 fold changes between sex comb and MS cells normalized by Z-score transformation (as in 35). Below each plot is a dot plot showing a selection of biological process terms found to be significantly enriched in each cluster relative to a background of all the genes detected in MS and comb shafts at one timepoint or more. (B-E) Time-series plots showing the proportion of reads that map to genes belonging to each of a selection of GO terms that we found to be significantly enriched in sex comb shafts. B-D are biological process terms; E is a cellular component term that encompasses both mitochondrial- and nuclear-encoded proteins localized to mitochondria. Vertical bars denote the bootstrapped confidence intervals around the median value, which is taken across all individuals cells se-quenced for a given cell type. (F) Adult male first tarsal segments from GAL4/UAS-mediated RNAi knockdown of electron transport chain and ATP synthase genes in the sex comb. subtypes of shaft, for genes that regulate several other aspects of filament formation, including polymerization factors, capping proteins, the Arp 2/3 complex, and proteins involved in vesicle-mediated monomer trafficking (**Supp. Fig. 14**).

The upregulated expression of the biochemical components of the ATP-producing machinery suggests that the morphological differences between sex comb teeth and MS bristles may in part arise from an increase in energy availability in the differentiating sex comb cells. To test this, we used the UAS-GAL4 system to reduce energy availability in the developing sex comb via the cell type-specific expression of RNAi constructs targeting electron transport chain and ATP synthase subunit genes. These perturbations led to clear changes in sex comb morphology, ranging from a reversion towards the shorter, thinner proportions of MS bristles when targeting components of ATP synthase (*ATP synthase subunits B* and *E*) to teeth that were markedly shorter even than MS bristles when targeting components of Mitochondrial Complex IV, the final ETC enzyme (*e.g., COX5B*) (**Fig. 4F**). These results suggest that differences in energy availability between sex comb and MS shafts are a major contributor to their morphological differences. Interestingly, however, the distinctive pigmentation of the comb teeth was largely unaffected, suggesting that pigmentation may be under the control of separate genetic modules or that the *dsx*-GAL4 we used is less active during the timeframe in which pigment production occurs.

### Sex comb shafts exhibit changes in the expression of genes that regulate actin filament depolymerization

The non-qualitative changes in gene expression we detect in sex comb cells relative to their MS homologs are well suited for increasing the rate of protein synthesis. The natural candidates for proteins being overproduced in the sex comb shafts are actins. After all, it’s the length of the actin bundles that ultimately determines shaft length (36). To our surprise, however, we detected no significant enrichment of actin-related GO terms in either the DPGP analysis or when running GO analyses on the sex comb shaft DEGs at each time point separately. When we looked at the expression profiles of individual actin-related genes, actin transcript abundance did not seem to correlate with the number of actin bundles known to be in the final bristle or with the overall volume of that cell (**Supp. Fig. 14**). Even more surprising was our observation that there was generally no clear difference in the expression of the bristle-expressed actins, *Act5C* or *Act42A* (37), between socket cells and the bristle-building shaft cells. The only exception was sex comb shaft-specific upregulation of *Act42A* at 36h APF, long after the size difference between sex comb teeth and MS shafts first becomes apparent (**Fig. 1G**).

Our failure to detect upregulation of actins in sex comb shafts may be a consequence of whole-transcriptome expansion brought about by endoreplication (see below), such that the absolute quantity of actin produced is increased while the proportion it represents of the total transcriptome remains unchanged. However, another explanation is that the amount of bundled actin is largely insensitive to the abundance of actin monomers *i.e.,* that actin is present in excess. In this case, the differences between longer and shorter bristles may be driven instead by factors that regulate the dynamics of actin assembly and disassembly. Consistent with this, we found persistent, cross-timepoint upregulation of the filament disassembly factors *tsr* (cofilin) and *flr* (AIP1) in sex comb shafts (**Supp. Fig. 14**). These factors prune and disassemble actin filaments that are not bundled or otherwise stabilized, a process of recycling that ensures the continued local availability of actin monomers at the tip, thereby facilitating the rapid growth of actin bundles (38). The upregulation of disassembly factors in the sex comb would therefore provide a mechanism to increase bristle growth rate. In contrast, for genes that regulate the bundling of actin filaments (*f, sn, jv*) we saw little compelling evidence of comb shaft upregulation, with the exception of slightly earlier upregulation of *sn* relative to MS shafts. As we observed for the bristle-expressed actins, we failed to see clear differences between shafts and sockets, let alone between

### Endoreplication contributes to the bristle-to-tooth transformation

The socket cells of the sex comb are also modified, being much larger than their MS homologs. To explore the genetic drivers of this transformation, we repeated the DPGP analysis on genes that were differentially expressed between comb and MS socket cells. There was considerable overlap with the results of the shaft analysis, with significant enrichment of terms relating to ATP production and protein synthesis **(Figure 5A; Supp. Fig. 15**). However, we also detected a cluster of genes with a sharp peak at 16h that was significantly enriched for terms relating to DNA replication. Given that we ran this analysis on the terminal socket cell populations, enrichment of DNA replication terms likely reflects differences in endoreplication rather than in the timing of mitosis. In principle, endoreplication offers a simple mechanism through which cell size can be increased by boosting the rate of protein synthesis. Existing data suggest that endoreplication is a normal part of bristle development. Indeed, bristle cells in different parts of the fly have been shown to differ in the number of endocycles they undergo, leading some to suggest that bristle size may be related to the degree of polyploidy (21, 23, 39).

**Figure 5.**
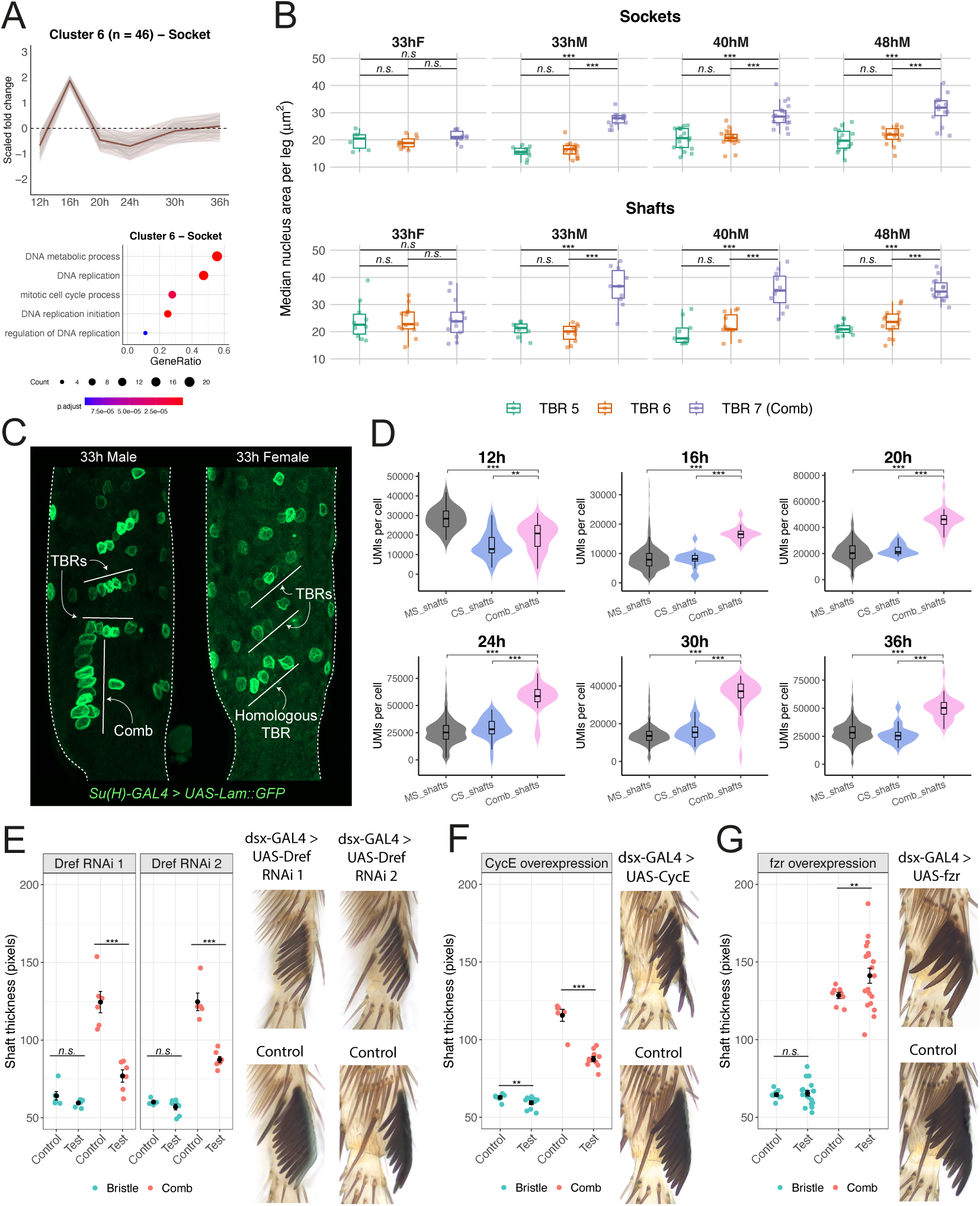
Increased endoreplication is a key part of the bristle-to-tooth transformation. (A) Expression trajectories for genes that were differentially expressed between sex comb and mechanosensory (MS) sockets and which were grouped into cluster 6 during the DPGP analysis. The data here are plotted using the same approach outlined in the Figure 4 legend. GO terms associat-ed with DNA replication that were significantly enriched in this cluster are shown below. The full set of cluster plots and associated GO terms are shown in Supp. Fig. 15). (B) Boxplots showing the median nucleus size (cross-section area) for each of the three most distal transverse bristle rows (TBRs) at three different timepoints in males (33h, 40h, and 48h after puparium formation, APF) and at one time point in females (33h APF). Data are plotted separately for the shaft and socket cells. Each individual dot represents a median value across bristles within a given row in a single fly at a given timepoint. p values were obtained by Tuk-ey-corrected contrasts between the estimated marginal means for each TBR/comb combination from a linear mixed effects model. (C) Representative images from a male and female at 33h APF expressing *UAS-Lamin::GFP* under the control of the *Su(H)*-GAL4 socket marker. White, undashed lines indicate the three most distal TBRs. In males, the most distal TBR is rotated 90 degrees and transformed into the sex comb. (D) The distribution of mRNA abundance (reads detected per cell) plotted separately for the different shaft cell populations and for each timepoint. A Kruskal-Wallis test was first used to determine if the effect of cell type on median read count was significant at a given timepoint. If so, pairwise Wilcoxon tests with Benjamini-Hochberg correction were then used to test for significant differences between each within-timepoint cell type pair. (E) Measurements of shaft thickness plot-ted separately for the MS bristles of the most distal non-comb TBR (blue) and comb teeth (red). In each panel, the ‘test’ genotype is *dsx*-GAL4 > *UAS-Dref RNAi*. Dref is a transcription factor known to regulate endoreplication in shaft cells (44). The panels differ only in the RNAi construct used in each case. RNAi 1 is TRiP.GLV21057 and RNAi 2 is TRiP.JF02232 (83). The control genotypes in each case are +/UAS males generated from the same cross. Each dot represents the mean taken across measured shafts for a single leg. The mean and standard error for each treatment combination is shown, as are representative images of the sex comb for each genotype. (F) As in (E) but for UAS-mediated overexpression of *CycE*. Overexpression of *CycE* has been shown to block endocycling (45–47). (G) As in (E) but for UAS-mediated overexpression of *fzr*, which triggers endoreplication in mitotic cells (48–50). n.s. denotes a non-statistically significant difference, ** = p<0.01, *** =p<0.001.

To test whether sex comb cells undergo additional endoreplication relative to MS bristles, we measured nuclear size—a proxy for DNA content (40–43)—of socket and shaft cells in the distal TBRs (TBRs 5 and 6) at multiple timepoints during the phase of active shaft growth. Not only was nuclear size significantly and consistently larger in comb sockets than in the TBRs, but it was significantly and consistently larger in comb shafts as well, suggesting that both cell types reach a higher ploidy level in the sex comb than in the surrounding MS bristles (**Fig. 5B,C**). Crucially, shaft and socket cells in the most distal TBR of the female first tarsal segment (the row homologous to the male sex comb) were not significantly larger than those in the adjacent TBRs, indicating that the elevated endocycling observed in their male homologs is a male-specific innovation rather than a positional effect (**Fig. 5B,C**). It’s unclear why we did not detect significant enrichment of DNA replication terms in our shaft DPGP analysis. It may be the result of temporal offsets between shafts and sockets in the timing of endoreplication, as is known to occur in the notum microchaetes (21). Further evidence for a ploidy increase in sex comb cells can be found in the scRNA-seq reads. From 16h APF, the number of reads recovered per cell in our scRNA-seq datasets was significantly higher for sex comb shafts than for MS or CS shafts (**Fig. 5D**). With the exception of 30h, when the difference was even greater, we persistently detected ∼2x as many reads in sex comb shafts than in MS (and CS) shaft cells (**Supp. Fig. 16A**). This closely parallels the ∼2x size difference we recorded between sex comb and MS shaft nuclei (**Fig. 5B**). Taken together, our data suggest that sex comb shafts undergo one additional round of endoreplication relative to their homologs in the TBRs.

To test whether ploidy differences contribute to the morphological differences between MS bristles and sex comb teeth, we used a *dsx*-GAL4 driver to knock down or overexpress genes implicated in the control of endoreplication. We began with efforts to reduce ploidy level in sex comb cells. RNAi-mediated knockdown of *Dref,* a transcription factor known to regulate endoreplication in shaft cells (44), led to a reduction in tooth thickness. Teeth from *Dref* knockdown males were intermediate in thickness between control teeth and MS bristles (**Fig. 5E**). Overexpression of *CycE,* which is tolerated in mitotically cycling cells but blocks endocycling (45–47), led to a similar reduction in tooth thickness (**Fig. 5F**). We next tried the opposite approach, attempting to boost ploidy level in sex comb cells. For this, we overexpressed *fzr,* which triggers endoreplication in mitotic cells (48–50) and which we found to be significantly upregulated in sex comb cells relative to MS bristles (**Supp. Fig. 16B**). Tooth number was reduced, likely due to some earlier divisions within the SOP lineage having been arrested, but average tooth thickness was significantly higher than in controls (**Fig. 5G**). Collectively, these experiments support a quantitative link between ploidy level and shaft size and suggest that increases in sex comb cell ploidy contribute to the morphological differences that distinguish sex comb teeth from their evolutionary progenitors, the MS bristles.

### The sex comb genetic pathway lacks clear modular architecture

Our data suggest that changes across a wide range of gene modules are required to transform a MS bristle into a sex comb tooth. At minimum, this includes ATP synthesis, protein synthesis, actin filament dynamics, and endoreplication. But how do these widespread gene expression changes connect back to the *dsx/Scr* ChIN that initiates the bristle-to-tooth transformation? Downstream of *dsx/Scr* we can envisage different regulatory architectures. That architecture may be modular, where *dsx/Scr* regulates the expression of several transcription factors, each of which then controls a separate gene network that controls one aspect of sex comb morphology (tooth size, curvature, pigmentation, *etc*.). Alternatively, the system may be highly decentralized, where each transcription factor contributes modestly or redundantly to multiple aspects of sex comb morphology. Our data are consistent with this latter model. We identified 17 transcription factors (excluding *dsx*) that were differentially expressed in sex comb shafts at any time point (**Supp. Fig. 17**). Of these, two (*fd96Ca* and *fd96Cb*) have previously been shown to have a modest effect on sex comb tooth morphology, with teeth slightly thinned on double knockdown under the control of *dsx*-GAL4 (51). For the remaining 15, we used *dsx*-GAL4 to knock down each in the developing sex comb. No individual knockdown had any effect on sex comb morphology. However, a double knockdown of *bab2,* which was one of the differentially expressed transcription factors, and its paralog *bab1*, which was not, induced a partial transformation of sex comb teeth towards the thinner, shorter morphology of the mechanosensory bristle, without affecting pigmentation (**Supp. Fig. 17**). Our data therefore argue against a modular architecture—no one transcription factor uncouples different aspects of sex comb morphology or is responsible for a total reversion to MS bristle morphology, and none of these transcription factors has a phenotype comparable to *dsx* or *Scr* loss of function (5, 52–54). Based on the *bab1/bab2* and *fd96Ca/ fd96Cb* knockdown phenotypes, it cannot be ruled out that these transcription factors positively regulate *Scr/dsx* levels and, by extension, that quantitative variation in *Scr/dsx* levels may generate quantitative variation in sex comb tooth morphology. If so, changes in *Scr/dsx* expression may provide a simple route through which some of the striking interspecific variation in sex comb tooth morphology (e.g., 5) has evolved.

## Discussion

The individualization of serially repeated homologs is a compelling model for the origin of evolutionary innovations: regulatory changes at the ChIN level decouple homologs from one another, downstream trait-building gene networks diverge, and new phenotypic endpoints are reached. Here, we characterized the gene expression divergence between the sex comb and MS bristles and showed that, instead of co-opting genes from other cell types or tissues, the newly evolved *dsx*-*Scr* ChIN has modified the timing and level of expression of genes that are present in the ancestral MS shaft transcriptome. The collective effect of these changes—which cover gene modules that regulate energy production, protein production, actin dynamics, and endoreplication—is to induce a sex-specific and spatially restricted shift in the tempo of bristle development, accelerating the rate of shaft growth to build something evolutionarily novel: a sex comb tooth (**Fig. 6**).

**Figure 6.**
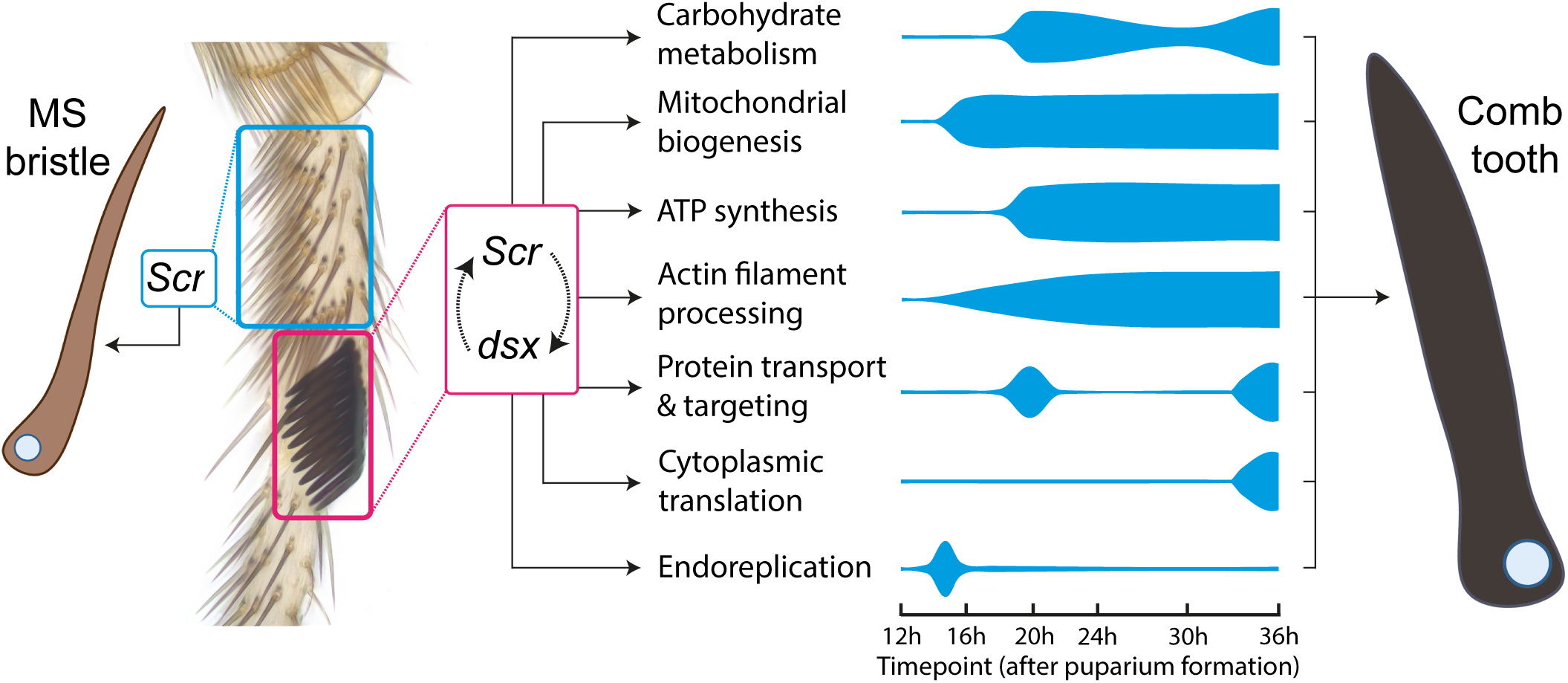
The bristle-to-tooth transformation is achieved through quantitative and heterochronic modulation of the ancestral mechanosensory bristle transcriptome. The gene programs that build the mechanosensory bristles of the transverse bristle rows (TBRs) in the first tarsal segment are downstream of a ChIN centered on the HOX gene *Scr* (shown in the blue square) (84, 85). In contrast, the ChIN that controls sex comb development centers on an autoregulatory feedback loop between *Scr* and the sex determination gene *dsx* (shown in the pink square) (28). Our data suggest that the *Scr-dsx* ChIN augments the expression of genes that are shared with mechanosensory bristle shafts, as opposed to activating the expression of genes that are absent from the mechanosensory bristle shafts. The collective effect of these expression changes is to accelerate the rate of shaft growth in sex comb teeth. The key biological processes that we find to be augmented are listed here in this figure, where the thickness of a ribbon denotes upregulation in sex comb shafts relative to mechanosensory bristles shafts. This representation is based on the timepoints at which we detected enrichment of related GO terms. The three exceptions to this are ‘mitochondrial biogenesis’, for which we use the data shown in Fig. 4E (the proportion of reads mapping to genes that are annotated with a cellular localization of ‘mitochondrion’); ‘actin filament processing’, for which we use the timepoints at which the actin filament disassembly factors *flr* (16h-36h APF) and *tsr* (24h-36h APF) were significantly upregulated (Supp. Fig. 14); and ‘endoreplication’, for which we use the comb and bristle read data shown in Fig. 5D (sex comb shafts only show their doubled read count, suggestive of elevated ploidy, from 16h APF onwards).

It is perhaps unsurprising that the larger size of sex comb teeth is associated with an increase in the energy-producing machinery. After all, a longer, thicker shaft requires more protein to build, and translation can account for >20-30% of total ATP consumption (55, 56), with the rate of protein synthesis highly sensitive to ATP levels (57). Higher mitochondrial mass and ATP availability correlate with higher levels of transcription and translation (58, 59), while experimentally increasing energy production can increase the rate of protein synthesis (60). This relationship between energy availability and protein production establishes metabolic regulation as a key determinant of developmental rate. Indeed, mutations that reduce mitochondrial mass slow down larval growth in *Drosophila* (61), and knockdown of electron transport chain components slows down development in *Caenorhabditis elegans* (62). Some of the best evidence for linkage between metabolic and developmental rates comes from the mammalian presomitic mesoderm. Oscillations of the segmentation clock are ∼2 times faster in mouse compared to human, correlating with higher mitochondrial density and higher metabolic rates in mouse cells (63–65). Reducing mitochondrial respiration rate inhibits protein synthesis and slows down the segmentation clock, although in this case the effect is mediated by NAD(H) redox homeostasis rather than by ATP supply (65). Similarly, the development of cortical neurons proceeds much slower in humans than mice (66), correlating with lower mitochondrial TCA cycle and electron transport chain activity in humans (67). Increasing oxidative metabolism in human neurons is sufficient to accelerate their morphological and functional maturation, while inhibiting mitochondrial activity in mouse neurons slows down their development (67). Our data build on these biochemical studies in two ways. First, by providing evidence that cell type-specific shifts in metabolic rate *within* an organism can drive between-organ variation in morphology. And second, by providing evidence that metabolically driven differences in developmental rate between cell types can have a strong transcriptional basis.

Alongside the upregulation of metabolic gene modules, our data implicate endoreplication as an additional mediator of developmental rate in bristle homologs. Our experiments showed that compromising endocycling in sex comb cells was associated with a reduction in tooth size and a reversion towards a more MS bristle-like morphology. A link between cell ploidy level and developmental rate seems intuitive: by amplifying gene copy number across the genome, endoreplication provides more transcriptional substrate, boosting the number of transcripts that can be produced per unit time—as evidenced by the elevated read count we recovered in comb shaft cells—and therefore the potential rate of protein synthesis. Consistent with this, cell ploidy level often correlates with cell size, suggesting that endoreplication provides a mechanism for increasing the rate or endpoint of cell growth (68). However, the relationship is not fixed, even within the same organism. In *Arabidopsis,* for example, polyploid cells grow at the same rate as diploid cells in the developing sepal (69), but much faster in the outer side of the apical hook (70). This discrepancy may be because an increase in nuclear DNA content is not itself directly responsible for stimulating an increase in cell growth; rather, cells undergo endoreplication in response to a perceived increase in growth rate in order to sustain continued growth (70–73). Recent work in *Arabidopsis* provides support for this, identifying a feedback loop where changes in the integrity of the cell wall—which is strained and thinned during growth—are used to control endoreplication and cell growth (70). Applying this logic to the sex comb, we could imagine that a shift in metabolic gene expression stimulates an increase in growth rate, which then feeds back to stimulate endoreplication to support that growth, perhaps via a mechanochemical signaling pathway centered on structural properties of the cell.

Increasing endocycle number and boosting metabolic gene expression both provide routes to increasing protein production. For the sex comb tooth, we might expect this increase to center on actins because it’s the number and length of bundled actin filaments that ultimately determine shaft size. Surprisingly, however, we do not observe major differences in the relative expression of actins and actin-associated genes. Membrane-tethered actin bundles form the structural framework of the bristle shaft, and can be visualized as parallel ridges in the shaft cuticle (25, 36). By this metric, sex comb teeth have more and longer actin bundles than most MS bristles, and yet the bristle actins *Act5C* and *Act42A* (37) do not show clear and persistent differential expression. The faster growth of sex comb teeth also indicates more rapid actin bundle assembly. While this rate is largely insensitive to the supply of actin or fascin (*singed*), which are present in excess, another actin crosslinking protein encoded by *forked* (*f*) is limited throughout bundle formation and regulates bundle size and shape (23, 74). However, *f* also does not show differential expression between comb teeth and MS bristle shafts. The most likely explanation for this result lies in the normalization of RNA counts data. Owing at least in part to their higher ploidy level, sex comb teeth contain many more transcripts than other bristle shafts (**Fig. 5D**). Thus, the absolute amounts of *Act5C* and *f* in sex comb teeth may be much higher than in MS shafts while their expression as a fraction of the overall transcriptome is approximately equal. The main actin-associated genes for which we see substantial expression differences are cofilin (*twinstar*) and its partner AIP1/*flare*, which are involved in the disassembly and renewal of actin filaments. In bristle shafts, highly cross-bridged and membrane-bound actin bundles turn over slowly and persist, whereas poorly crosslinked filaments turn over more rapidly and disappear (23). Cofilin promotes the growth of actin bundles by maintaining actin monomer supply and by disassembling unbundled filaments and branched networks that compete with stabilized bundles (38), suggesting that elevated expression of *tsr* and *flr* may contribute to the faster growth of actin bundles in sex comb teeth.

Our data provide a counterpoint to the growing body of work that implicates gene co-option as a central driver of evolutionary innovation. Gene co-option is clearly an important evolutionary force (75), as a wide body of examples from the pipefish’s broodpouch (76) to various novelties seen across drosophilid genitalia (8, 10) attest to. Indeed, the recurrent, independent redeployment of wing-building genes alone has been associated with such varied morphological innovations as the *Tribolium castaneum* ‘gin-trap’ (77), the leafhopper helmet (78), and the prothoracic horns of scarabaeine beetles (79). However, while gene co-option may be a common route through which tissues, organs, and cell types diverge from one another (80)—perhaps even the general route—our data show that the co-option of genes into a trait-building gene network is not necessary and that quantitative and heterochronic shifts in the expression of a conserved gene set can drive evolutionary innovation. The capacity to innovate at the morphological level may be particularly potent when ChINs evolve to modulate the expression of genes that control metabolic rate, as our data suggest occurs in the sex comb. This potency emerges in part from the ability to exploit existing dynamic links between metabolic rate and aspects of cellular physiology, such as those that might link growth rate to endoreplication (70). Where such links are present, a simple regulatory change that affects metabolic rate would trigger a cascade of downstream changes that could collectively support a morphological transformation. Metabolically driven differences in developmental rate may therefore provide a powerful alternative to gene co-option as an evolutionary driver of divergence between serial homologs.

## Supplementary figure legends

**Supplementary figure 1. Sex comb cells differentiate earlier than mechanosensory bristle cells.** Stainings of an 18h APF male first tarsal segment with antibodies against Pax2 and Su(H). Whereas Fig. 1D prioritizes the view of the sex comb, this image prioritizes the view of the transverse bristle rows (TBRs). CS=chemosensory bristles.

**Supplementary figure 2. Shaft growth is accelerated in the sex comb relative to the surrounding mechanosensory bristles.** (A) The output of a likelihood ratio test comparing two linear mixed effects models that model the effect of organ ‘class’ (sex comb or mechanosensory bristle) and developmental stage (‘Age’: 32h, 40h, and 48h after puparium formation) on shaft length. One model includes the interaction term while the other omits it. As random effects, we include the three length measurements taken per individual bristle (‘BristleID’) nested within an individual leg (‘LegID’). The significant *p*-value (*p* <0.0001) indicates that the inclusion of the interaction term significantly improves model fit and, therefore, that the difference in shaft length between the sex comb teeth and mechanosensory (MS) bristles increases with age. (B) F-values and p-values from the full model specified in (A), *i.e.,* the interaction term model. (C) Estimated marginal means and confidence limits derived from the full model using the R package ‘emmeans’. (D) The estimated marginal means plotted in (C). (E) The difference in estimated marginal mean lengths between teeth in the comb (*i.e.,* TBR 7) and MS bristles (TBR 6). (F) Within-timepoint contrasts conducted on the emmeans output using the function ‘pairs’ and a Tukey *p*-value adjustment. The contrast estimates are plotted in (E) and the *p-*values correspond to those plotted in (C).

**Supplementary figure 3. Identification of sensory cell populations in the 12h APF male dataset.** (A) A UMAP of the 12h male dataset with major cell populations annotated. Annotation nomenclature follows that used in Hopkins *et al.* (81). (B) Genes enriched for expression in each annotated cluster labelled in (A). (C) The sensory organ lineage cells shown in (A), subsetted out and reclustered with cell types annotated. (D) Genes enriched for expression in each annotated cluster labelled in (C). Note that this dataset was generated from *dsx*-GAL4 > *UAS-mCherry* individuals, hence the inclusion of *mCherry* in the dotplot. MS=Mechanosensory, CS=Chemosensory, Camp.=Campaniform sensillum.

**Supplementary figure 4. Identification of sensory cell populations in the 16h APF male dataset.** (A) A UMAP of the 16h male dataset with major cell populations annotated. Annotation nomenclature follows that used in Hopkins *et al.* (81). (B) Genes enriched for expression in each annotated cluster labelled in (A). (C) The sensory organ lineage cells shown in (A), subsetted out and reclustered with cell types annotated. (D) Genes enriched for expression in each annotated cluster labelled in (C). Note that this dataset was generated from *dsx*-GAL4 > *UAS-GFP* individuals, hence the inclusion of *GFP* in the dotplot. MS=Mechanosensory, CS=Chemosensory, Camp.=Campaniform sensillum.

**Supplementary figure 5. Identification of sensory cell populations in the 20h APF male dataset.** (A) A UMAP of the 20h male dataset with major cell populations annotated. Annotation nomenclature follows that used in Hopkins *et al.* (81). (B) Genes enriched for expression in each annotated cluster labelled in (A). (C) The sensory organ lineage cells shown in (A), subsetted out and reclustered with cell types annotated. (D) Genes enriched for expression in each annotated cluster labelled in (C). Note that this dataset was generated from *dsx*-GAL4 > *UAS-mCherry* individuals, hence the inclusion of *mCherry* in the dotplot. MS=Mechanosensory, CS=Chemosensory.

**Supplementary figure 6. Identification of sensory cell populations in the 24h APF male dataset.** (A) A UMAP of the 24h male dataset with major cell populations annotated. Annotation nomenclature follows that used in Hopkins *et al.* (81). (B) Genes enriched for expression in each annotated cluster labelled in (A). (C) The sensory organ lineage cells shown in (A), subsetted out and reclustered with cell types annotated. (D) Genes enriched for expression in each annotated cluster labelled in (C). Note that this dataset was generated from *RAL-517* individuals, so no fluorescent markers are shown. An orange bar denotes subpopulations of chemosensory neuron that were readily identifiable. *nvy^+^*and *fkh*^+^ neurons follow the nomenclature used in Hopkins *et al.* (2023). PS neurons are the pheromone-sensing neurons, which at this clustering resolution could not readily be separated into the male- and female-sensing cells seen in Hopkins *et al.* MS=Mechanosensory, CS=Chemosensory.

**Supplementary figure 7. Identification of sensory cell populations in the 30h APF male dataset.** (A) A UMAP of the 30h male dataset with major cell populations annotated. Annotation nomenclature follows that used in Hopkins *et al.* (81). (B) Genes enriched for expression in each annotated cluster labelled in (A). (C) The sensory organ lineage cells shown in (A), subsetted out and reclustered with cell types annotated. (D) Genes enriched for expression in each annotated cluster labelled in (C). Note that this dataset was generated from *RAL-517* individuals, so no fluorescent markers are shown. An orange bar denotes subpopulations of chemosensory neuron that were readily identifiable. *nvy^+^*and *fkh*^+^ neurons follow the nomenclature used in Hopkins *et al.* (2023). PS neurons are the pheromone-sensing neurons, which at this clustering resolution could not readily be separated into the male- and female-sensing cells seen in Hopkins *et al.* MS=Mechanosensory, CS=Chemosensory, Camp.=Campaniform sensillum.

**Supplementary figure 8. Identification of sensory cell populations in the 36h APF male dataset.** (A) A UMAP of the 36h male dataset with major cell populations annotated. Annotation nomenclature follows that used in Hopkins *et al.* (81) with the new addition of trichome cells, which express *ovo* and *sha*. Wr. glia = Wrapping glia. Sur. glia = Surface glia. (B) Genes enriched for expression in each annotated cluster labelled in (A). (C) The sensory organ lineage cells shown in (A), subsetted out and reclustered with cell types annotated. (D) Genes enriched for expression in each annotated cluster labelled in (C). Note that this dataset was generated from *dsx*-GAL4 > *UAS-mCherry* individuals, hence the inclusion of *mCherry* in the dotplot. An orange bar denotes subpopulations of chemosensory neuron that were readily identifiable. *nvy^+^, fkh*^+^, male pheromone-sensing (MPS) and female pheromone-sensing (FPS) neurons follow the nomenclature used in Hopkins *et al.* MS=Mechanosensory, CS=Chemosensory, Camp.=Campaniform sensillum.

**Supplementary figure 9. The distinct clustering of sex comb cells is not an artifact of the expression of *Gal4 and mCherry.*** A UMAP of the sensory cells in the 12h APF male dataset after removing *Gal4* and *mCherry* and reclustering. Cells are coloured with the annotation they were given in the dataset that includes *Gal4* and *mCherry.* The comb socket and shaft populations still form distinct clusters indicating that their separation is a product of natural differences in expression. Arrows denote the differentiation trajectories. pIIIb cells divide to form neurons and sheaths, while pIIa cells divide to form sockets and shafts.

**Supplementary figure 10. Accessory cells in chemosensory bristles express genes that are largely absent from their mechanosensory and sex comb homologues.** (A-C) HCR *in situ* stainings against genes that are heavily enriched in chemosensory (CS) bristles relative to homologous cells in other bristle classes. *CG32040, mtg,* and *CG14280* in shafts, *Ance-3* in sockets, and *NLaz* in sheaths. We use *Su(H)* as a socket marker and *acj6* as a marker of 3 of the 4 neurons that innervate each CS bristle (81). The *mtg* probe shows high background staining throughout the leg but localized staining in the CS shaft is clear. Note that *mtg* staining in (A) is shown only at 27h APF and *acj6* staining in (C) is shown only at 27h APF and 40h APF. Asterisks denote the position of readily identifiable CS bristles in (B) and (C). (D) Time-series expression plots for genes largely specific to CS shaft and socket cells.

**Supplementary figure 11. *black* is specific to the sex comb in the first tarsal segment but is not required for normal sex comb development.** (A) HCR in situ staining of *b* and *pros* at 30h APF in a male and female first tarsal segment. *pros* is used here as a bristle marker, being expressed in the sheath cell of every bristle and the neurons of chemosensory bristles (81). The position of the sex comb is labeled, as is the likely position of the pre-apical (‘pre-ap’) bristle in the distal tibia. (B) First tarsal segments from adult males that are transheterozygous for mutant alleles affecting *b*. These alleles are a molecular null (*b[1]*)(82), a hypomorphic GAL4 line (*b-GAL4*)(86), and a deficiency that covers the *b* locus on chromosome arm 2L (*Df(2L)*). Genotypes heterozygous for the molecular null or deficiency line over either the CyO or SM6a balancer are shown as controls. The pre-apical bristle in the distal tibia is labelled in one image. Note how transheterozygotes show the darker cuticular melanization phenotype known to be associated with *b* nulls (82) but no obvious defects in the sex comb.

**Supplementary figure 12. Quantitative upregulation of genes in the sex comb and chemosensory bristles.** Time-series expression plots for genes showing significant quantitative expression differences between sex comb and mechanosensory bristle cells, without evidence of heterochrony. The exception is *aqz*, which shows quantitative upregulation in chemosensory rather than sex comb cells.

**Supplementary figure 13. Expression trajectories for genes found to be differentially expressed between sex comb shafts and their mechanosensory homologs.** All clusters that are not present in Figure 4 are shown. Genes are grouped based on the cluster they were assigned to in our DPGP analysis. Each expression plot shows the trajectory of each gene in the cluster (gray lines) as well as the cluster mean (thick, colored line). The ribbon denotes ± 2 S.D. The number of genes in the cluster is given in the plot title. The values plotted are the log2 fold changes between sex comb and mechanosensory (MS) cells normalized by Z-score transformation (as in 35). Below each plot is a dot plot showing a selection of up to 5 biological process GO terms found to be significantly enriched in each cluster relative to a background of all the genes detected in MS and comb shaft cells at one timepoint or more.

**Supplementary figure 14. Sex comb shafts modulate gene networks that regulate actin filament depolymerization.** Time-series gene expression plots, grouped by the actin-related function each gene performs. Sex comb-enriched expression of the filament disassembly factors *flr* and *tsr* (highlighted with red boxes) provide a potential route through which shaft growth can be accelerated. By pruning and disassembling filaments that are not bundled, these factors ensure the continued availability of actin monomers at the growing tip (38). Note that in many cases, there is no clear difference between shafts and sockets, let alone different classes of shaft.

**Supplementary figure 15. Expression trajectories for genes found to be differentially expressed between sex comb sockets and their mechanosensory homologs.** All clusters that are not present in Fig. 5 are shown. Genes are grouped based on the cluster they were assigned to in our DPGP analysis. Each expression plot shows the trajectory of each gene in the cluster (gray lines) as well as the cluster mean (thick, colored line). The ribbon denotes ± 2 S.D. The number of genes in the cluster is given in the plot title. The values plotted are the log2 fold changes between sex comb and mechanosensory (MS) cells normalized by Z-score transformation (as in 35). Below each plot is a dot plot showing a selection of up to 5 biological process terms found to be significantly enriched in each cluster relative to a background of all the genes detected in MS and comb socket cells at one timepoint or more.

**Supplementary figure 16. Changes in endoreplication dynamics are a key part of the bristle-to-tooth transformation.** (A) The ratio of sex comb shaft to mechanosensory (MS) shaft and chemosensory (CS) shaft read counts at each timepoint. A value of 1 suggests equivalent read counts between, for instance, MS shaft and comb shaft cells, while a value of 2 means that we detect twice as many reads in comb cells. Bootstrapped confidence intervals (2000 iterations) for the ratios are presented. (B) Time-series expression plots for endoreplication-mediating genes we tested for roles in sex comb development using the UAS/GAL4 system. Asterisks denote timepoints where a gene was significantly upregulated in sex comb shafts relative to mechanosensory shafts. Of the three genes we tested—*Dref, CycE,* and *fzr*—only *fzr* was significantly upregulated in sex comb cells relative to MS bristles at any time point in our scRNA-seq data, raising the possibility that it might be a direct regulator of the ploidy differences between sex comb and mechanosensory cells. However, we did not see a corresponding increase in MS bristles when overexpressing *fzr* under the control of the bristle driver *Pax2-*GAL4, suggesting that *fzr* alone is insufficient to drive the additional endocycles seen in sex comb cells.

**Supplementary figure 17. Transcription factor knockdowns in the developing sex comb suggest a decentralized genetic pathway.** Expression plots for each transcription factor that we detected as significantly upregulated in sex comb cells at any timepoint. With the exception of *dsx* and the forkhead domain genes *fd96Ca* and *fd96Cb* (which have been tested previously, 51), we used UAS-RNAi constructs to knockdown the expression of each transcription factor in the developing sex comb under the control of *dsx-*GAL4. Only a double knockdown of *bab1* and *bab2* (87) gave a phenotype, showing an incomplete transformation of sex comb teeth towards the thinner, shorter proportions of a mechanosensory bristle.

**Supplementary figure 18. Processing the single-cell data.** (A) The number of predicted cells recovered in each dataset when running CellRanger without ‘force-cells’, alongside the final number of cells in each dataset after filtering cells out based on dataset-specific quality control thresholds. (B) The contamination fraction estimated in the 20h APF dataset when forcing CellRanger to detect set numbers of cells. The dashed red line indicates the number of cells detected (15,911) using default settings (*i.e.,* without force-cells). (C) The distribution of UMIs detected per barcode in the 20h dataset when using default settings or forcing the detection of 12,000 cells. Note the reduction in low barcode ‘cells’ (*i.e.,* putative empty droplets) when instructing CellRanger to force the detection of fewer cells. (D) The distribution of UMIs detected per barcode in each dataset when forcing the detection of 12,000 cells. Note that the female 24h dataset also includes the distribution when forcing the detection at 14,000 cells. We used a higher value for this dataset because cutting at 12,000 lost some of the lower part of the distribution that was retained in all other datasets. (E) A UMAP of the full, force-cells, 20h dataset without any downstream filtering of cells based on quality control metrics. Note how low UMI cells generally form their own cluster or fall between closely associated clusters in UMAP space (*e.g.,* between chemosensory and mechanosensory neuron clusters), suggestive of their aberrant characteristics. (F) The distribution of UMIs, features, and percentage of reads mapping to mitochondrial genes for each of the datasets pre- and post-filtering based on cell-level quality control metrics. The post-filtering median value is provided as a label in each case.

**Supplementary table 1.** The top 30 genes contributing to the putative epithelial-contamination gene expression program (GEP) in each dataset, derived from a Consensus Non-negative Matrix Factorization (cNMF) analysis.

**Supplementary table 2.** The proportion of reads that the putative epithelial-contamination GEP accounts for in each cell. Cells are binned according to the proportion of the transcriptome that this GEP accounted for in each dataset. Cells with values greater than 10% were removed from our non-epithelial cell datasets on the basis that they are likely doublets that include epithelial cells. The proportion of cells removed from each dataset is also provided in this table.

## Materials & Methods

### Fly strains and husbandry

Flies used in all experiments were raised on a standard cornmeal medium and housed in an incubator at 25°C on a 12:12 cycle. For the 24h and 30h APF single-cell RNA-seq samples that we previously published (81), we used individuals from the DGRP wildtype strain *RAL-517* (88). For the 16h APF sample we used progeny from a cross between *dsx-GAL4* (89) and *UAS-GFP.nls* (BDSC: #4775). For the 12h, 20h, and 36h APF samples, we used progeny from a cross between *dsx-GAL4* and *UAS-mCherry.nls* (BDSC: #38424). For visualizing shaft length, we crossed *neur-GAL4* (BDSC: #6393) to *UAS-Act5C::RFP* (BDSC: #24778). For visualizing nuclear area, we used *UAS-Lam::GFP* (BDSC: #7376) with *Pax2-GAL4* (a gift from Joshua Kavaler) for shafts and *Su(H)-GAL4.ASE4* for sockets (BDSC: #93029). Other lines used: *UAS-COX5A RNAi* (BDSC: #27548)*, UAS-ATPsynB RNAi* (BDSC: #27712)*, UAS-ATPsynE RNAi* (BDSC: #67826)*, b[1]* (BDSC: #227), Mi{Trojan-GAL4.0}b[MI10547-TG4.0]/SM6a (BDSC: #76724), Df(2L)FDD-0428643/SM6a (BDSC: #25166), Df(2L)BSC252/CyO (BDSC: #23152), *UAS-bab1 siRNA #3 bab2 siRNA #16* (87), *UAS-fzr* (BDSC: #91689)*, UAS-CycE.L* (BDSC: #4781), *UAS-CrebA RNAi* (BDSC: #31900 and #27548)*, UAS-CG3328 RNAi* (BDSC: #55211 and #57713)*, UAS-CG5446 RNAi* (BDSC: #40824)*, UAS-CG17002 RNAi* (BDSC: #82969)*, UAS-CG4374 RNAi* (BDSC: #57745 and #61252)*, UAS-COX5B RNAi* (BDSC: #80383), *UAS-cwo RNAi* (BDSC: #26318 and #27736)*, UAS-MED17 RNAi* (BDSC: #34664)*, UAS-mid RNAi* (BDSC: #50681)*, UAS-Nf-YA RNAi* (BDSC: #63542)*, UAS-Pdp1 RNAi* (BDSC: #26212)*, UAS-Sp1 RNAi* (BDSC: #35777)*, UAS-svp RNAi* (BDSC: #44394)*, UAS-tx RNAi* (BDSC: #25973 and #34792), and *UAS-Dref RNAi* (BDSC: #31941 and #35692).

### Single-cell suspension preparation

We prepared our single-cell suspensions from dissected first tarsal segment tissue following the protocol we previously developed as part of our cell atlas (81) and which is available on protocols.io (90). Briefly, white prepupae were collected, sexed, and transferred to folded kimwipe wet with water (300μl for 12h and 16h, 450μl for 20h, and 750μl for 36h) held inside a petri dish. The dish was moved to an incubator at 25°C on a 12:12 cycle and the pupae left to age. Pupae were removed from puparia using forceps and placed on top of a water-soaked kimwipe 1h before the desired age was reached (*e.g.,* 23h after collection for the 24h sample). Pupae were then placed ventral side up on tape, the base of the abdomen pierced to release fluid pressure, and the foreleg removed at the tibia/tarsal joint. The dissected leg was moved to a piece of tape, covered in a drop of 1X Dulbecco’s PBS (DPBS, Sigma, D8537), and a Micro Knife (Fine Science Tools, 10318–14) used to sever at the midpoint of the second tarsal segment. The first tarsal segment was then eased out of the pupal cuticle and transferred to a glass well on ice containing 100μl of 1X DPBS using a BSA-coated 10μl tip.

Once 65-70 segments were collected, the DPBS was removed from the well and replaced with 100μl of dissociation buffer, which consisted of 10X TrypLE (Thermo Fisher Scientific, A12177-01) with a final concentration of 2 mg/mL of collagenase (Sigma, C0130). The well was sealed and submerged in a metal bead bath in an incubator at 37°C for 35 minutes. For the 36h APF sample, we incubated for 45 minutes to overcome the greater resistance to dissociation at this point. The dissociation buffer was then removed from the well and replaced with 50μl of room temperature DPBS before the suspension was pipetted up and down 20-40x using a 200μl, widebore, low bind, freshly BSA coated tip (Thermo Fisher Scientific, 2069G) on a 50μl pipette set to 40μl. The solution was then pipetted up and down a further 20-40x using a flame-rounded, BSA-coated 200μl tip, again on a 50μl pipette set to 40μl. Next, the solution was transferred to a 2mL, low-bind, wide-bottomed tube on ice. Cell concentration and viability was assayed using an acridine orange/propidium iodide stain and measured using a LUNA-FL Fluorescent Cell Counter averaged across 2 × 5μl aliquots.

### Generation of tarsal single-cell data

Barcoded 3’ single cell libraries were prepared from single cell suspensions using the Chromium Next GEM Single Cell 3’ kit v3.1(10X Genomics, Pleasanton, California) for sequencing according to the manufacturer’s instructions. The cDNA and library fragment size distribution were verified on a Bioanalyzer 2100 (Agilent, Santa Clara, CA). The libraries were quantified by fluorometry on a Qubit instrument (LifeTechnologies, Carlsbad, CA) and by qPCR with a Kapa Library Quant kit (Kapa Biosystems-Roche) prior to sequencing. Libraries from the 16h, 20h, 24h, 30h, and 36h APF samples were sequenced on a NovaSeq 6000 sequencer (Illumina, San Diego, CA) with paired-end 150 bp reads. The 12h APF libraries were sequenced in a 28 x 10 x 10 x 90 bp configuration on an AVITI sequencer (Element Biosciences, San Diego, CA). In each case, the sequencing generated approximately 50,000 reads per cell and 500 million reads per library.

### Processing of single-cell data

The reference index was built using the ‘mkref’ function in CellRanger (v7.0.0) and the Ensembl BDGP6.32 *Drosophila melanogaster* genome into which we introduced sequences for GAL4, GFP, and mCherry. The GTF was filtered to include features of gene_biotype ‘ncRNA’ and ‘protein_coding’. Alignment, barcode assignment, and UMI counting were performed on each single-cell dataset using the ‘count’ function in CellRanger. For each dataset, we ran ‘count’ with default settings (including counting regions that mapped to intronic regions), other than using the ‘force-cells’ option to limit the number of cells detected to 12,000 (except in the case of the 24h female dataset where we used 14,000; see below). We opted to use ‘force-cells’ because across our datasets, CellRanger’s cell detection algorithm routinely detected more than the 10,000 cells we expected to recover based on the volume of cell suspension loaded (**Supp. Fig. 18A**). A consequence of including the full complement of cells identified by CellRanger (*i.e.,* without use of ‘force-cells’) was, in some datasets, a marked increase in the degree of ambient RNA contamination that we estimated using SoupX (91). For example, the contamination fraction was estimated at 0.64 when using the full 15,911 cells identified by CellRanger in the 20h dataset, a value that likely represents a failure in estimating the contamination fraction (as suggested by the SoupX documentation). Consistent with this, the estimated contamination fraction fell to 0.08 when forcing CellRanger to call either 10,000, 11,000, or 12,000 cells (**Supp. Fig. 18B**), which falls within the 0% to 10% range most commonly seen for single cell (rather than nuclei) experiments (91).

A substantial proportion of cells above the 10,000-cell mark are likely to be empty droplets containing ambient RNA or droplets containing damaged cells. This conclusion is supported by the bimodal distribution that we observed in the number of UMIs detected per barcode (**Supp. Fig. 18C**). The peak at lower UMI-per-barcode values was reduced, but not lost, when overriding CellRanger’s cell identification calls to force the selection of 12,000 cells (**Supp. Fig. 18C**). At this 12,000-cell cutoff, the bimodality persisted in all but the 24h female dataset, leading us to force CellRanger to detect 14,000 cells in this dataset (**Supp. Fig. 18D**). To further filter out suspect cells, we used dataset-specific UMI thresholds to remove cells falling within the lower peak. The cells filtered out during this process generally formed their own cluster and were often observed to fall between closely associated clusters in UMAP space (*e.g.,* between CS and MS neuron clusters), reinforcing their suspect characteristics (**Supp. Fig. 18E**). We further removed cells in each dataset in which fewer than 450 genes were detected and where >10% of reads mapped to mitochondrial genes. Our filtered, SoupX-corrected datasets ranged from including 8851 to 13487 cells (**Supp. Fig. 18A**), with a median value of 5352 to 12614 UMIs, 1238 to 2233 genes, and 0.56% to 0.93% reads mapping to mitochondrial genes per cell (**Supp. Fig. 18F**).

Each individual filtered, SoupX-corrected dataset was then run through a standard processing pipeline we built using Seurat (v4.4.0) (92–95). Data were normalized using “SCTransform”, regressing out variation due to the percentage of reads mapping to mitochondrial reads and selecting 5000 variable features. “RunPCA” (npcs=100), “FindNeighbors” (dims=1:100), “FindClusters” (resolution=2.2), and “RunUMAP” (dims=1:100) were each then run in succession. Using the resulting UMAP and following our previously published methods (81), we were able to readily identify and remove epithelial cells, which formed a contiguous cluster that accounted for the majority of cells in each dataset. From the new non-epithelial dataset, we then reapplied gene-level filtering to retain only those genes expressed in 3 or more cells and reran SCTransform and the same downstream clustering pipeline, this time using npcs=50, dims=50, and resolution=0.7.

As in our previously published cell atlas, many of the doublets identified by DoubletFinder (96) seemed suspicious when run on the full dataset and when run on the non-epithelial subsets in some of our newer datasets, for example heavily targeting *stg*^+^ mitotic neurons. We therefore sought alternative approaches to identify putative doublets. To this end, we used Consensus Non-negative Matrix Factorization (cNMF), a program designed to infer gene expression programs from scRNA-seq data (GEP)(97). Our idea was that doublets should show anomalous combinations of GEPs. Running cNMF (max-nmf-iter 2000, -k 8-40, -niter 250) and using the k_selection_plot to identify the optimal, dataset-specific choice of k, we observed that in each dataset, we detected a GEP enriched for a panel of genes that showed no cluster-specificity, were widely distributed across clusters, and enriched in epithelial cells when visualized in epithelial cell-containing datasets. The identities of these genes were largely shared between this GEP in each of our datasets (Supplementary table 1). We calculated the proportion of reads that this putative epithelial GEP accounted for in each cell and removed cells with values greater than 10% on the basis that they were likely doublets that included epithelial cells, which are the most common cell type in the dataset. This generally led to the removal of between 2.1% and 11.5% of cells from the non-epithelial datasets, although the figure was higher in the female 24h dataset, where 18.5% of cells were removed (Supplementary table 2). With putative epithelial doublets removed, we then reran the gene-level filtering and clustering pipeline, using the same values described previously. We then used the resulting UMAP to annotate non-epithelial cell types based on the panel of cell type-specific markers we previously published (81).

### Differential expression, clustering, and gene ontology analyses

To identify differentially expressed genes (DEGs), we first log-normalized SoupX-corrected RNA counts using the NormalizeData function with default parameters. We then used a Wilcoxon rank sum test implemented through the ‘FindMarkers’ function in Seurat, comparing either sex comb cells or CS cells to MS cells. We ran this separately for shaft and socket cells. For each analysis, we returned only positive markers and specified that, in order to be tested, a gene had to be expressed in a minimum of 10% of cells in either of the two populations being compared and to exceed a log fold change of 0.25. A Bonferroni correction was applied to p-values and an adjusted p-value cut off-of 0.05 used to determine significance. For time-series plots of individual gene expression, we used the Seurat function ‘AverageExpression’ to estimate the average, log normalized expression of a gene across cells in a given cluster at a given timepoint.

We used DPGP (35) to cluster genes that were differentially expressed between sex comb and MS cells in relation to their temporal trajectories. Unlike other clustering approaches that use Euclidean distance or correlation, such as k-mean and hierarchical clustering, DPGP accounts for the non-independence of consecutive timepoints and is therefore specifically designed to handle time-series data (35). We used log2 fold changes as the input into our DPGP analyses, performing separate analyses for shafts and sockets. We ran DPGP with default parameters, except for defining a low alpha value (0.001) to minimize the number of returned clusters. We built custom ggplot2 (v3.4.4) scripts to plot the trajectories of genes in each cluster returned by the DPGP analysis. For these plots, we applied a Z-score transformation to the log2 fold change values, subtracting the cluster mean from each expression value and dividing by the cluster standard deviation (as in 35). To test for significant enrichment of functional classes among the genes in each cluster, we used a gene ontology analysis implemented through the R package clusterProfiler (v4.14.6). We specified ‘biological process’ as the ontology classification, ‘Benjamini-Hochberg’ to adjust p-values, a p-value cut-off of 0.05, and q-value cut-off of 0.2. As a background gene set for the comparison, we used all genes expressed in at least 3 cells at at least one timepoint across all MS and comb cells for sockets and shafts, separately. For the organism database we used ‘org.Dm.eg.db’ (v3.20.0). We included a selection of up to 5 terms in summary dot plots, using the function ‘simplify’ to reduce redundancy among enriched GO terms (cutoff = 0.8, by = qvalue). For a subset of GO terms, we extracted all genes annotated to a given GO term and its descendant terms using the AnnotationDbi package. For each cell, we then calculated the sum of counts across all genes in that set and divided it by the total number of counts. For each cluster, we then calculate the median proportion across cells and used the boot function in the R package ‘boot’ (v1.3-28.1) to calculate bootstrapped confidence intervals using 1000 replicates.

### Fixation, immunohistochemistry, and microscopy

White P1 prepupae (0 to 1 h APF) were collected and sexed under light microscope. Pupae were then placed on a damp kimwipe and aged in an incubator at 25°C until the desired time point. Pupae were then placed on sticky tape and a razor blade used to cut away the dorsal half. The cut pupae were fixed in a 4% paraformaldehyde solution (125 μl 32% PFA, 675 μl H_2_O, 200 μl 5X TN) for 50 min on a rotator at room temperature and stored at 4°C. For leg dissections, fixed pupae were removed from the puparia in 1X TNT and tears made in the pupal cuticle at the femur–tibia boundary using forceps. The tibia through to ta5 was then pulled through the tear, freeing it from the pupal cuticle. At this point, legs expressing fluorescent proteins were mounted in Fluoromount 50 (SouthernBiotech). For immunohistochemistry, the dissected leg region was blocked with 5% goat serum (200μl 10% goat serum with 200μl 1X TNT) overnight at 4°C. Legs were then incubated with primary antibody solution overnight at 4°C, washed for 21 mins 4 times in 1X TNT, incubated with the secondary antibody solution for 2h at room temperature, washed for 21 mins 4 times in 1X TNT, and then mounted in Fluoromount 50 (SouthernBiotech). All stages, from dissection through to staining, were carried out in a glass well. Antibodies were used in solution with 1X TNT and a final concentration of 2% goat serum. Anti-Pax2 rabbit (a gift from Joshua Kavaler) was used at a final concentration of 1:5000, anti-Su(H) mouse at 1:200, anti-mouse AlexaFluor594 (Thermo Fisher Scientific Cat# A-11005, RRID:AB_2534073) at 1:400, and anti-Rabbit AlexaFluor488 (Thermo Fisher Scientific Cat# A-11008, RRID:AB_143165) at 1:400. Confocal images were taken using a Zeiss 980 Airyscan. Image stacks were processed using Z-series projection in ImageJ.

### Hybridization chain reaction *in situ*

For HCR *in situs,* we followed a lightly modified version of the HCR v3.0 protocol published by Bruce *et al.* (98), which is itself a modified version of Choi *et al.*’s (99) original HCR v3.0 protocol. Pupal legs fixed following the procedures outlined in the previous section were first washed three times (10min, 5min, 5min) in PTw (1X PBS with 0.1% Tween 20) in a glass well on a rotator. We then permeabilized the tissue in 200μl of detergent solution for 30min at room temperature before replacing the solution with 200μl of pre-warmed probe hybridization buffer. For 40mL of probe hybridization buffer, we mixed 12mL formamide, 10mL of 20X sodium chloride sodium citrate (SSC), 360μl 1 M citric acid pH 6.0, 400μl of 10% Tween 20, 200μl of 10 mg/mL heparin, 800μl of 50X Denhardt’s solution, 8 mL of 50% dextran sulfate, and filled up to 40 mL with distilled H_2_O. We then removed the hybridization buffer and replaced with probe solution, which was 200μl of the hybridization buffer with 2μl (from 1μM stock solution) of each probe. Tissues were then incubated overnight at 37C without rotation. The probe solution was then removed and washed 4x 15min with 200μl of pre-warmed probe washer buffer at 37C on a rotator. To make 40 mL of the probe wash buffer, we followed the procedure for the probe hybridization buffer but left out the Denhardt’s solution and dextran sulfate. The tissue was then washed a further 2x 5min with 200μl of 5X SSCT (5X SSC and 0.1% Tween 20) at room temperature on a rotator before replacing with 200μl of pre-equilibrated amplification buffer for 30min at room temperature. To make 40mL of amplification buffer, we mixed 10 mL of 20X SSC, 400μl of 10% Tween 20, 8 mL of 50% dextran sulfate, and filled up to 40 mL with distilled H_2_O. The amplification buffer was then removed and replaced with the hairpin solution—which was 200μl of the amplification buffer into which we added 2μl of hairpin 1 and 2μl of hairpin 2 for each probe used—and incubated overnight in the dark at room temperature. Prior to use, this hairpin buffer had been heated to 95C for 90 seconds before cooling to room temperature in a dark drawer for 30min. The tissue was then washed 5x (1×5min, 1×5min, 1×30min, 1×30min, 1×5min) with 200μl of SSCT and mounted in fluoromount.

B1 488, B2 594, and B3 647 Hairpins were ordered from Molecular Instruments, Inc. (Los Angeles, CA, USA). Probes were ordered as oligopools from Integrated DNA Technologies, Inc., (Coralville, Iowa, USA). In some cases, these probe pools targeted one gene and in other cases we combined probes targeting 3 different genes into a single pool. 50pmol probes received from IDT were diluted in 50ul of elution buffer (Qiagen) and stored at −20C in 10ul aliquots. Probes were designed using previously published software (https://github.com/rwnull/HCRProbeMakerCL)(100) and with the following settings: between 30-70% CG bases in a probe, a maximum A/T homopolymer length of 5, and a maximum C/G homopolymer length of 3.

### Nuclei area measurements and analysis

To visualize nuclei, we expressed a GFP-tagged form of the nuclear envelope protein Lamin (CG6944) under the control of a socket (*Su(H)-*GAL4) or shaft (*Pax2*-GAL4) driver. Because the GFP signal was too weak earlier in development, we focused our measurements of nuclear area on 3 time points—33h, 40h, and 48h APF—that spanned much of the period of active shaft growth in both the TBRs and sex comb. We reasoned that further increases in ploidy were unlikely to occur following the end of shaft growth, a judgement supported both by endoreplication timing in the notum (21) and by the fact that both the comb and TBR shafts are secreting cuticle by 48h APF, thereby preventing further shaft growth. Consequently, any further endoreplication beyond 48h APF seems unlikely.

Nuclear area was measured manually by using the freehand selection tool in ImageJ to trace the outline of each nucleus three times. To test for differences in nuclear area between bristle rows (including the comb) and timepoints, we constructed and analyzed separate linear mixed effects models for, separately, the male socket and shaft raw data using the R packages lme4 (v1.1-35.1) and lmerTest (v0.9-40). For both cell types, the model structure was NuclearArea ∼ BristleRow*Timepoint + (1|LegID) + (1|MeasurementNumber), *i.e.,* separate measurements of each nuclei and the ID of the leg were included as random effects. For sockets, we measured 986 nuclei across 47 legs and found significant effects of age (F_2,44.94_ = 8.77, p < 0.001), bristle row (TBR 5, TBR 6, comb; F_2,937.61_ = 634.37, p < 0.001), and the interaction between them (F_4,937.08_ = 4.08, p = 0.003). For shafts, we measured 707 nuclei across 35 legs and found significant effects of bristle row (F_2,670.71_ = 504.44, p < 0.001) and the interaction between class and age (F_4,670.56_ = 4.75, p < 0.001), but not age individually (F_2,32.06_ = 0.54, p = 0.587). *p*-values for the fixed effects were calculated using Satterthwaite’s method. For the pairwise, posthoc comparisons, we used the emmeans package (v1.10.0) to compute estimated marginal means for the interaction term (BristleRow*Timepoint) in the full model and performed comparisons between all factor levels. *p*-values from these comparisons were corrected using the Tukey method. This same statistical pipeline was used to test for sex effects in a dataset of all 33h nuclei across both males and females. Here, the model was NuclearArea ∼ BristleRow*Sex + (1|LegID) + (1|MeasurementNumber). For both sockets and shafts we found significant effects of bristle row (socket: F_2,367.25_ = 94.25, p < 0.001; shaft: F_2,317.85_ = 107.81, p < 0.001) and the interaction between row and sex (socket: F_2,367.25_ = 61.84, p < 0.001; shaft: F_2,317.85_ = 83.01, p < 0.001), but not sex individually (socket: F_1,27.37_ = 0.97, p = 0.33; shaft: F_1,20.63_ = 0.16, p = 0.690). For the plots presented in the text, a median value was taken for each nuclei across the three measurements, and then a median across nuclei for each bristle row was taken.

### Shaft measurements and analysis

Shaft length in developing pupae was measured from images of *neur-GAL4 > UAS-Act5c::RFP* males using the segmented line tool in ImageJ. Measurements were taken from the distal tip of the shaft and traced proximally along its length to the base. Shaft thickness in adult legs was measured at the base of the bristle/tooth. Three measurements were taken for each unobscured bristle/tooth in the sex comb and distal TBR. Analyses were performed using an analogous statistical framework to that used for nuclei, *i.e.* model comparison between linear mixed effects model followed by post-hoc comparisons implemented via emmeans.

## Data availability

Raw sequencing reads and preprocessed sequence data are available through NCBI GEO with accession code GSE293356. All other code and datasets used in this paper, including processed Seurat objects with cell type annotations, are available on the Open Science Framework at https://osf.io/25bnc.

## Acknowledgements.

We thank Hong Qiu for preparing libraires from our single-cell suspensions and performing the sequencing. We also thank Ryal Null and Rachel Thayer for advice on designing HCR probes and running the *in situs,* Joshua Kavaler for providing anti-Pax2 antibodies, Shyama Nandakumar for sharing her thoughts on endoreplication with us, the Bloomington *Drosophila* Stock Center for strains, the Developmental Studies Hybridoma Bank for antibodies, and Flybase for providing an indispensable source of information on gene function. We acknowledge the use of BioRender in designing elements used in our figures. Our graphical abstract incorporates a modified drawing of *D. melanogaster* from SciDraw (https://doi.org/10.5281/zenodo.3925939). This study was made possible in part through access to the MCB Light Microscopy Core Facility, the NIH grant S10OD026702 that allowed funding of the Zeiss 980 LSM, and training and support by Thomas Wilkop. This work was supported by NIH grant R35 GM122592 to A.K., a long-term fellowship from the Human Frontier Science Program Organization (LT000123/2020-L) to B.R.H., and start-up funds from the University of Florida to B.R.H.

## Author contributions

Conceptualization, B.R.H. and A.K.; Data curation, B.R.H.; Formal analysis, B.R.H.; Funding acquisition, B.R.H. and A.K.; Investigation, B.R.H., O.B., X.W., M.M.S, H.A.B., and S.N.; Methodology, B.R.H., O.B., and A.K.; Project administration, B.R.H. and A.K.; Software, B.R.H.; Project supervision, B.R.H. and A.K.; Visualization, B.R.H.; Writing, B.R.H. and A.K.

## Notes

https://www.ncbi.nlm.nih.gov/geo/query/acc.cgi?acc=GSE293356

https://osf.io/25bnc

### Competing Interest Statement

The authors have declared no competing interest.

