## Supplementary figures for "Morphological innovation without gene co-option: the *Drosophila* sex comb evolved via changes in developmental tempo and energy metabolism"

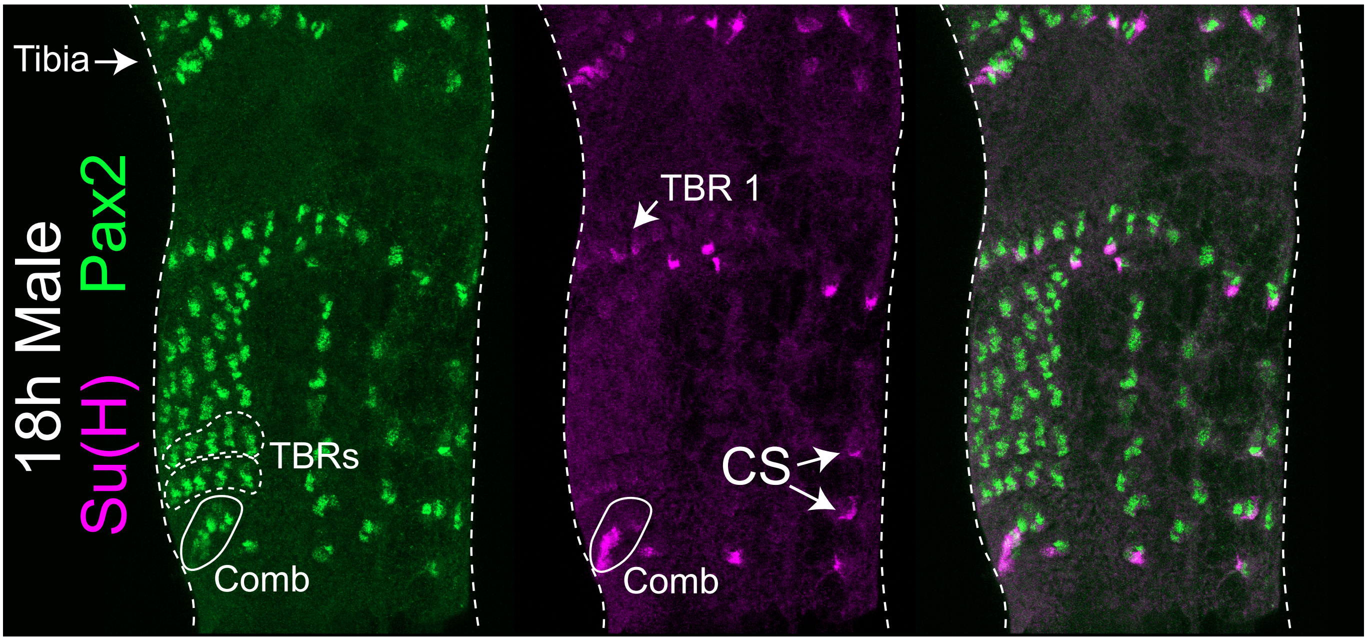


**Supplementary figure 1. Sex comb cells differentiate earlier than mechanosensory bristle cells.** Stainings of an 18h APF male first tarsal segment with antibodies against Pax2 and Su(H). Whereas Fig. 1D prioritizes the view of the sex comb, this image prioritizes the view of the transverse bristle rows (TBRs). CS=chemosensory bristles.

A

| Model Comparison: Likelihood Ratio Test | | | | | | | | |
|---|---|---|---|---|---|---|---|---|
| Model | npar | AIC | BIC | logLik | deviance | Chisq | Df | Pr(>Chisq) |
| Length ~ Class + Age + (1 \| LegID/BristleID) | 7.000 | 3798.151 | 3836.770 | −1892.075 | 3784.151 | | | |
| Length ~ Class * Age + (1 \| LegID/BristleID) | 9.000 | 3750.542 | 3800.195 | −1866.271 | 3732.542 | 51.609 | 2.000 | <0.0001 |

B

| Type III Analysis of Variance Table with Satterthwaite's Method | | | | | | |
|---|---|---|---|---|---|---|
| Fixed effects | Sum Sq | Mean Sq | NumDF | DenDF | F value | Pr(>F) |
| Class | 127.272 | 127.272 | 1.000 | 587.222 | 1154.546 | <0.0001 |
| Age | 65.635 | 32.817 | 2.000 | 68.731 | 297.702 | <0.0001 |
| Class:Age | 5.892 | 2.946 | 2.000 | 587.191 | 26.725 | <0.0001 |

C

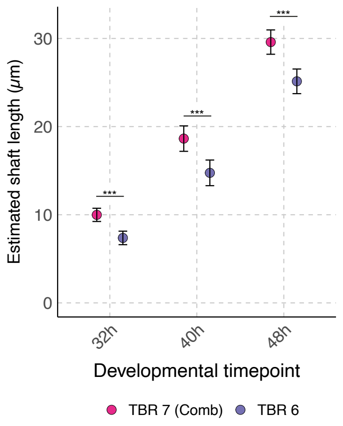


E

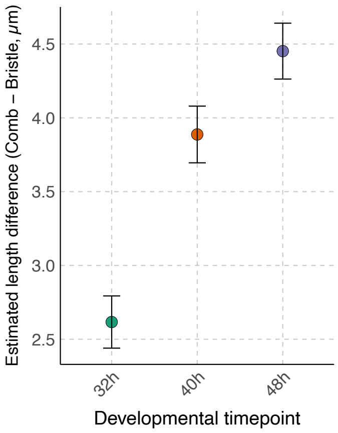


D

| Estimated Shaft Lengths by Age and Bristle Row | | | | | |
|---|---|---|---|---|---|
| | Est. mean | SE | df | Lower CL | Upper CL |
| 32h | | | | | |
| TBR 7 (Comb) | 9.987 | 0.377 | 81.113 | 9.237 | 10.737 |
| TBR 6 | 7.370 | 0.386 | 89.177 | 6.603 | 8.137 |
| 40h | | | | | |
| TBR 7 (Comb) | 18.640 | 0.721 | 56.376 | 17.197 | 20.084 |
| TBR 6 | 14.753 | 0.727 | 58.543 | 13.297 | 16.209 |
| 48h | | | | | |
| TBR 7 (Comb) | 29.589 | 0.689 | 56.213 | 28.208 | 30.970 |
| TBR 6 | 25.137 | 0.698 | 59.209 | 23.740 | 26.535 |

F

| Model Estimates | | | | | | |
|---|---|---|---|---|---|---|
| contrast | Age | estimate | SE | df | t.ratio | p.value |
| TBR 7 (Comb) - TBR 6 | 32h | 2.617 | 0.177 | 605.787 | 14.789 | <0.0001 |
| TBR 7 (Comb) - TBR 6 | 40h | 3.887 | 0.192 | 570.030 | 20.239 | <0.0001 |
| TBR 7 (Comb) - TBR 6 | 48h | 4.452 | 0.189 | 570.052 | 23.520 | <0.0001 |

**Supplementary figure 2. Shaft growth is accelerated in the sex comb relative to the surrounding mechanosensory bristles.** (A) The output of a likelihood ratio test comparing two linear mixed effects models that model the effect of organ 'class' (sex comb or mechanosensory bristle) and developmental stage ('Age': 32h, 40h, and 48h after puparium formation) on shaft length. One model includes the interaction term while the other omits it. As random effects, we include the three length measurements taken per individual bristle ('BristleID') nested within an individual leg ('LegID'). The significant p-value ($p < 0.0001$) indicates that the inclusion of the interaction term significantly improves model fit and, therefore, that the difference in shaft length between the sex comb teeth and mechanosensory (MS) bristles increases with age. (B) F-values and p-values from the full model specified in (A), i.e., the interaction term model. (C) Estimated marginal means and confidence limits derived from the full model using the R package 'emmeans'. (D) The estimated marginal means plotted in (C). (E) The difference in estimated marginal mean lengths between teeth in the comb (i.e., TBR 7) and MS bristles (TBR 6). (F) Within-timepoint contrasts conducted on the emmeans output using the function 'pairs' and a Tukey p-value adjustment. The contrast estimates are plotted in (E) and the p-values correspond to those plotted in (C).

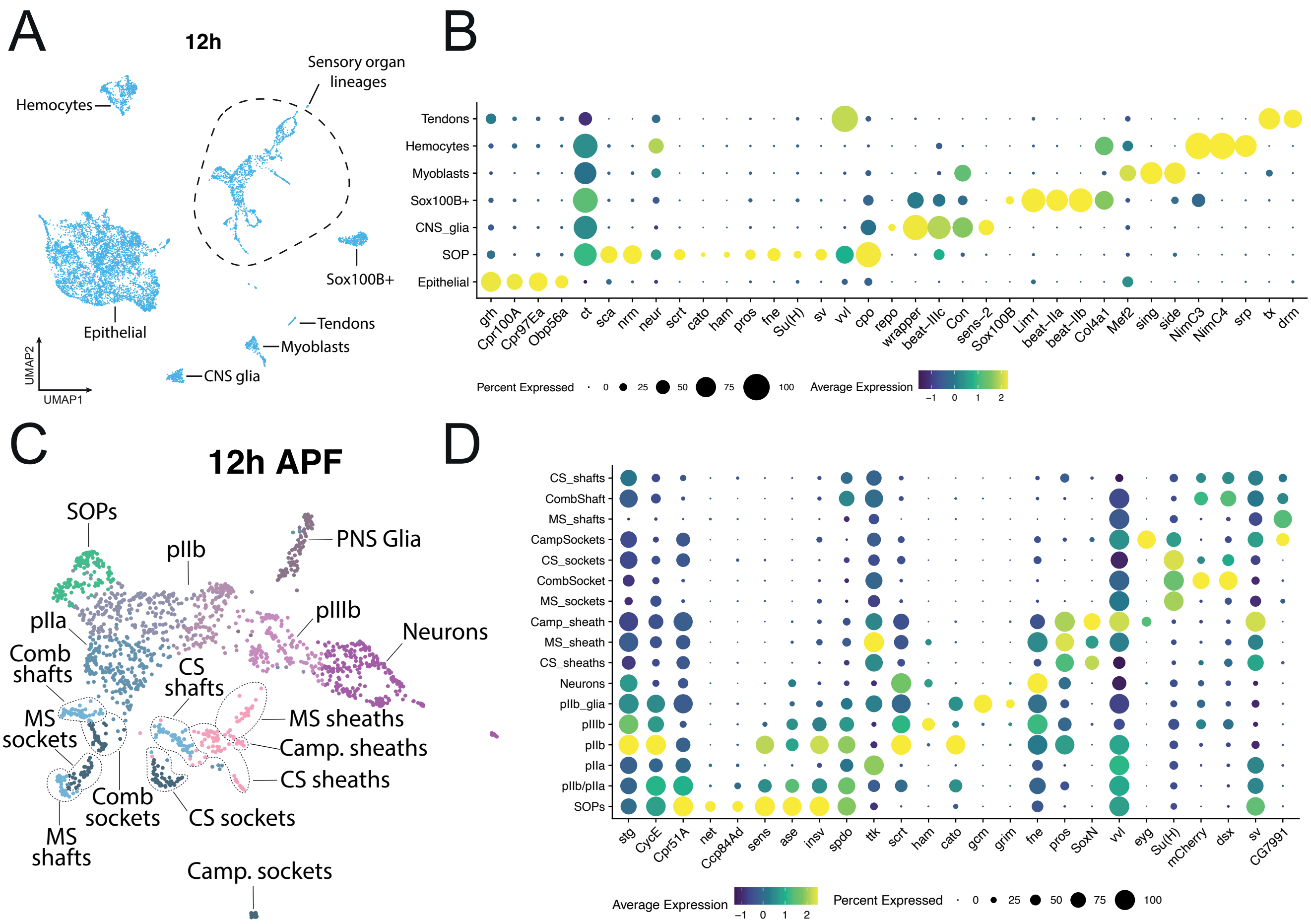


**Supplementary figure 3. Identification of sensory cell populations in the 12h APF male dataset.** (A) A UMAP of the 12h male dataset with major cell populations annotated. Annotation nomenclature follows that used in Hopkins *et al.* (81). (B) Genes enriched for expression in each annotated cluster labelled in (A). (C) The sensory organ lineage cells shown in (A), subsetted out and reclustered with cell types annotated. (D) Genes enriched for expression in each annotated cluster labelled in (C). Note that this dataset was generated from *dsx*-GAL4 > *UAS-mCherry* individuals, hence the inclusion of *mCherry* in the dotplot. MS=Mechanosensory, CS=Chemosensory, Camp.=Campaniform sensillum.

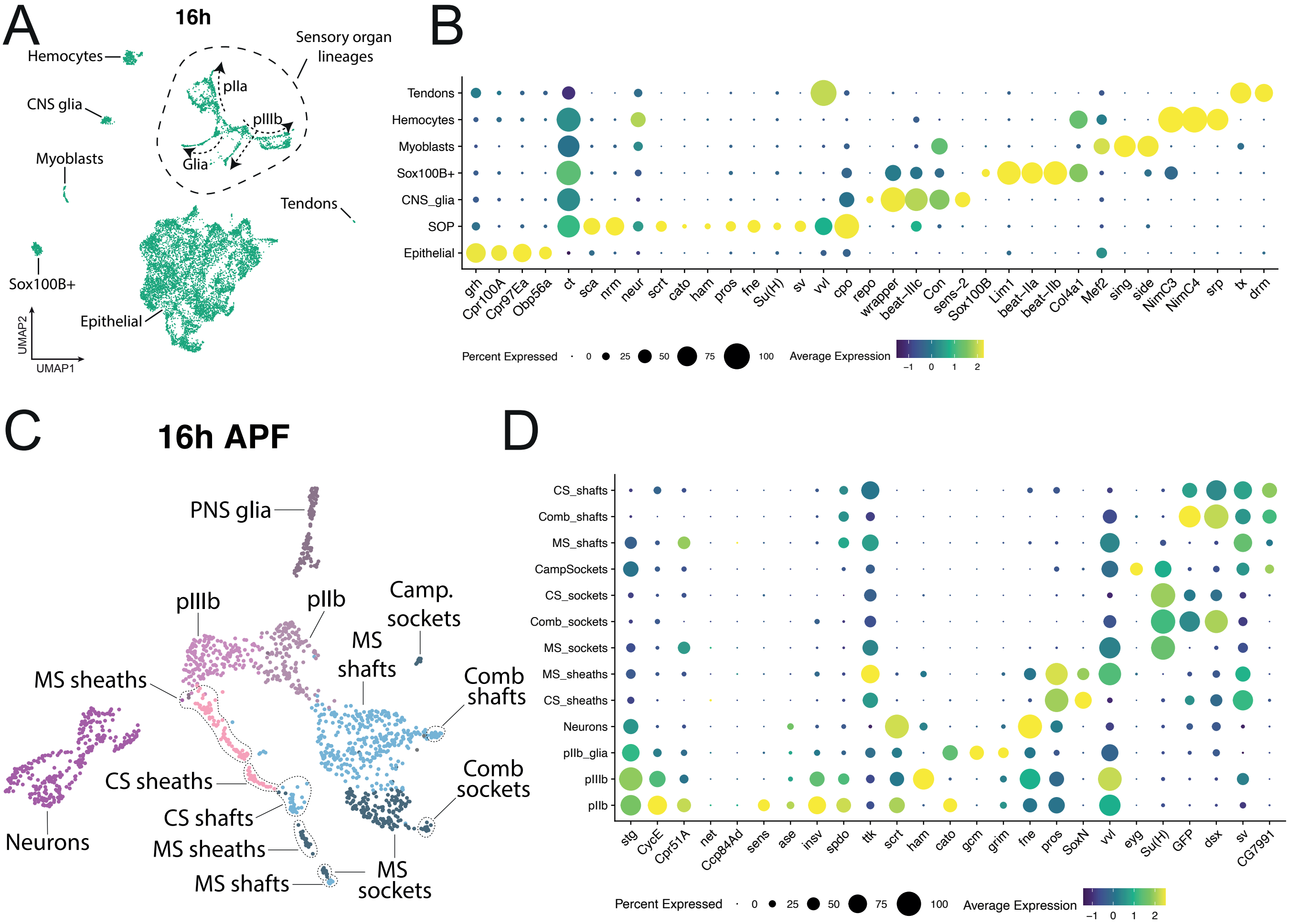


**Supplementary figure 4. Identification of sensory cell populations in the 16h APF male dataset.** (A) A UMAP of the 16h male dataset with major cell populations annotated. Annotation nomenclature follows that used in Hopkins *et al.* (81). (B) Genes enriched for expression in each annotated cluster labelled in (A). (C) The sensory organ lineage cells shown in (A), subsetted out and reclustered with cell types annotated. (D) Genes enriched for expression in each annotated cluster labelled in (C). Note that this dataset was generated from *dsx*-GAL4 > *UAS-GFP* individuals, hence the inclusion of *GFP* in the dotplot. MS=Mechanosensory, CS=Chemosensory, Camp.=Campaniform sensillum.

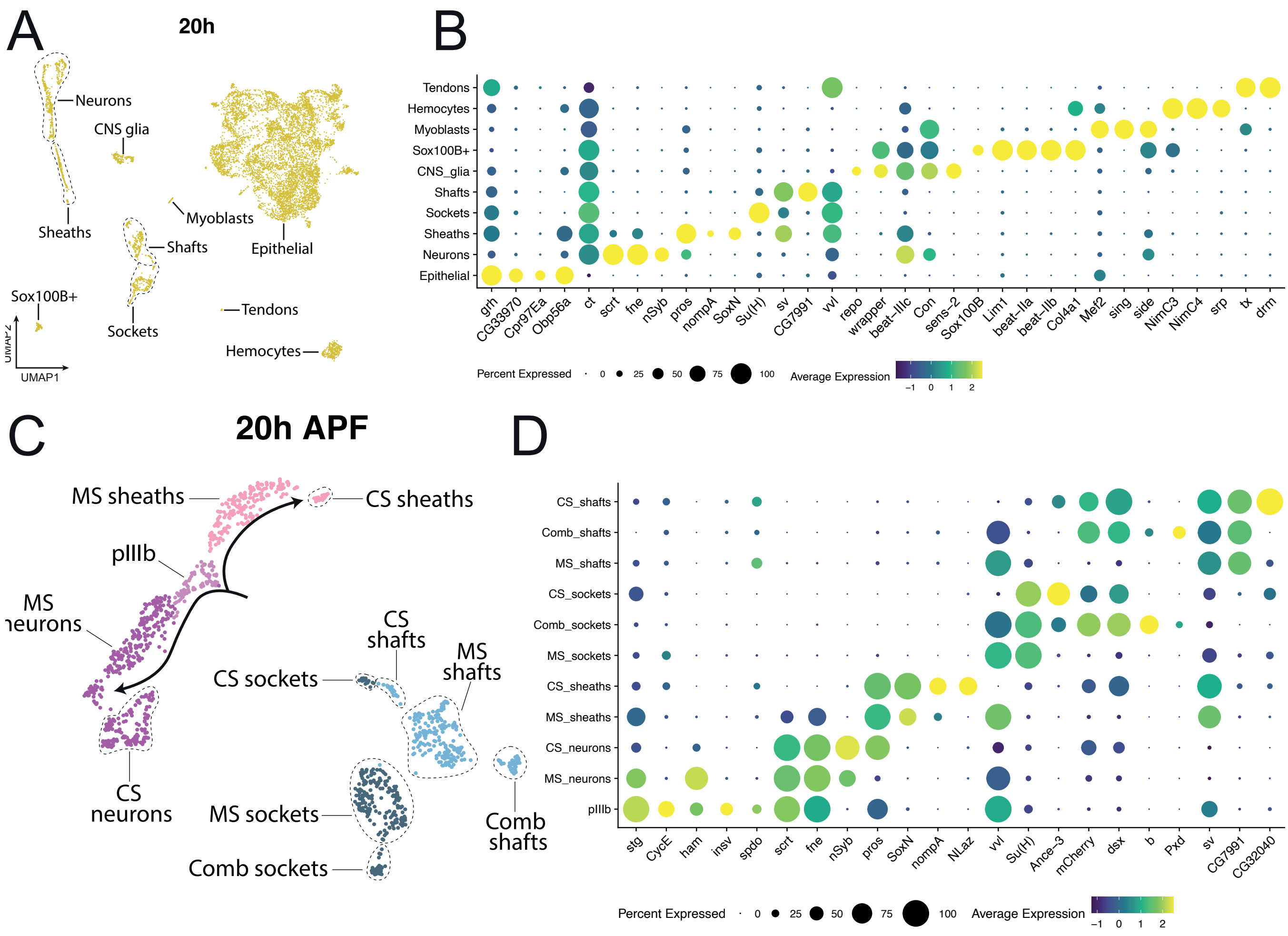


**Supplementary figure 5. Identification of sensory cell populations in the 20h APF male dataset.** (A) A UMAP of the 20h male dataset with major cell populations annotated. Annotation nomenclature follows that used in Hopkins *et al.* (81). (B) Genes enriched for expression in each annotated cluster labelled in (A). (C) The sensory organ lineage cells shown in (A), subsetted out and reclustered with cell types annotated. (D) Genes enriched for expression in each annotated cluster labelled in (C). Note that this dataset was generated from *dsx*-GAL4 > *UAS-mCherry* individuals, hence the inclusion of *mCherry* in the dotplot. MS=Mechanosensory, CS=Chemosensory.

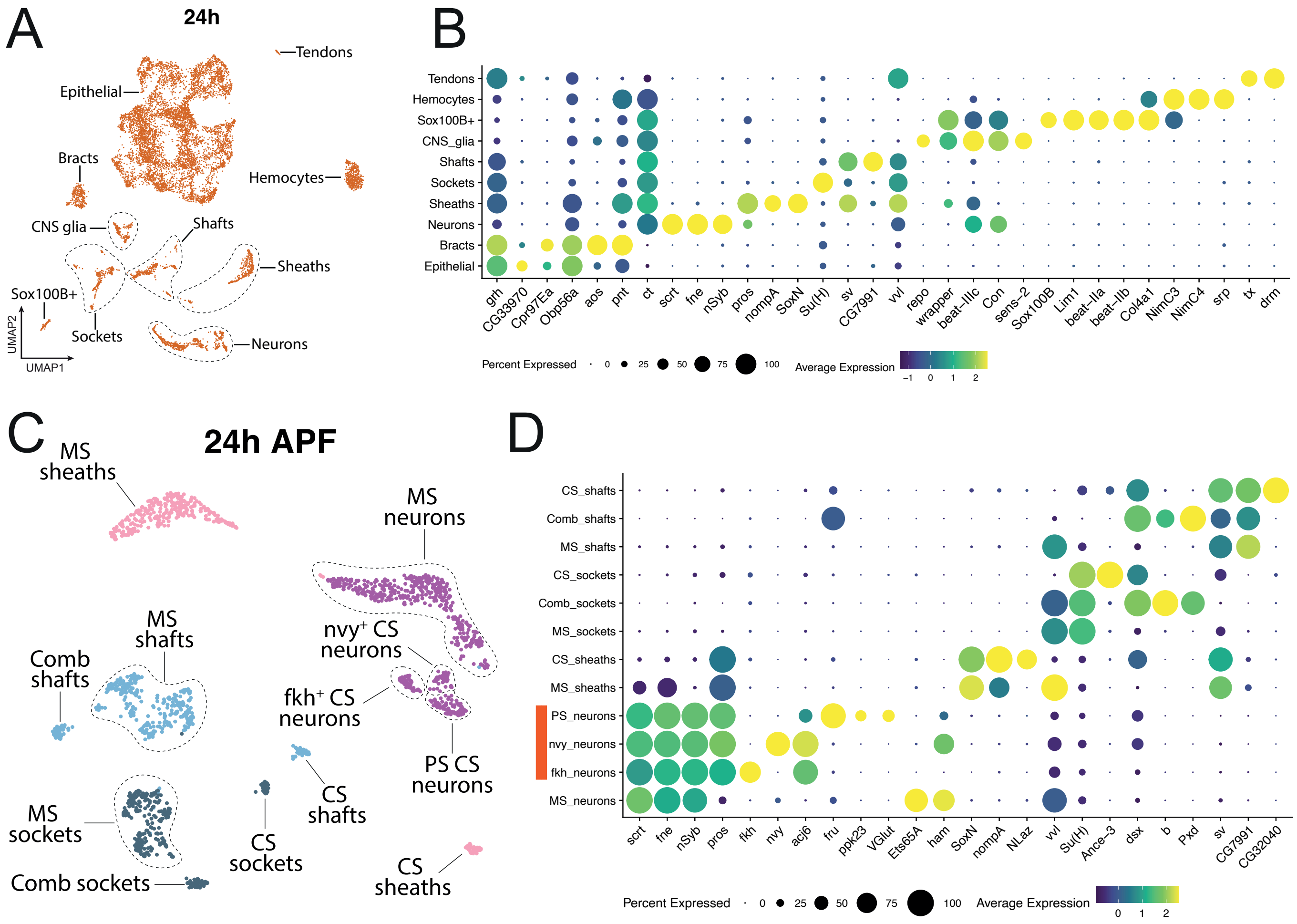


**Supplementary figure 6. Identification of sensory cell populations in the 24h APF male dataset.** (A) A UMAP of the 24h male dataset with major cell populations annotated. Annotation nomenclature follows that used in Hopkins *et al.* (81). (B) Genes enriched for expression in each annotated cluster labelled in (A). (C) The sensory organ lineage cells shown in (A), subsetted out and reclustered with cell types annotated. (D) Genes enriched for expression in each annotated cluster labelled in (C). Note that this dataset was generated from *RAL-517* individuals, so no fluorescent markers are shown. An orange bar denotes subpopulations of chemosensory neuron that were readily identifiable. nvy+ and fkh+ neurons follow the nomenclature used in Hopkins *et al.* (2023). PS neurons are the pheromone-sensing neurons, which at this clustering resolution could not readily be separated into the male- and female-sensing cells seen in Hopkins *et al.* MS=Mechanosensory, CS=Chemosensory.

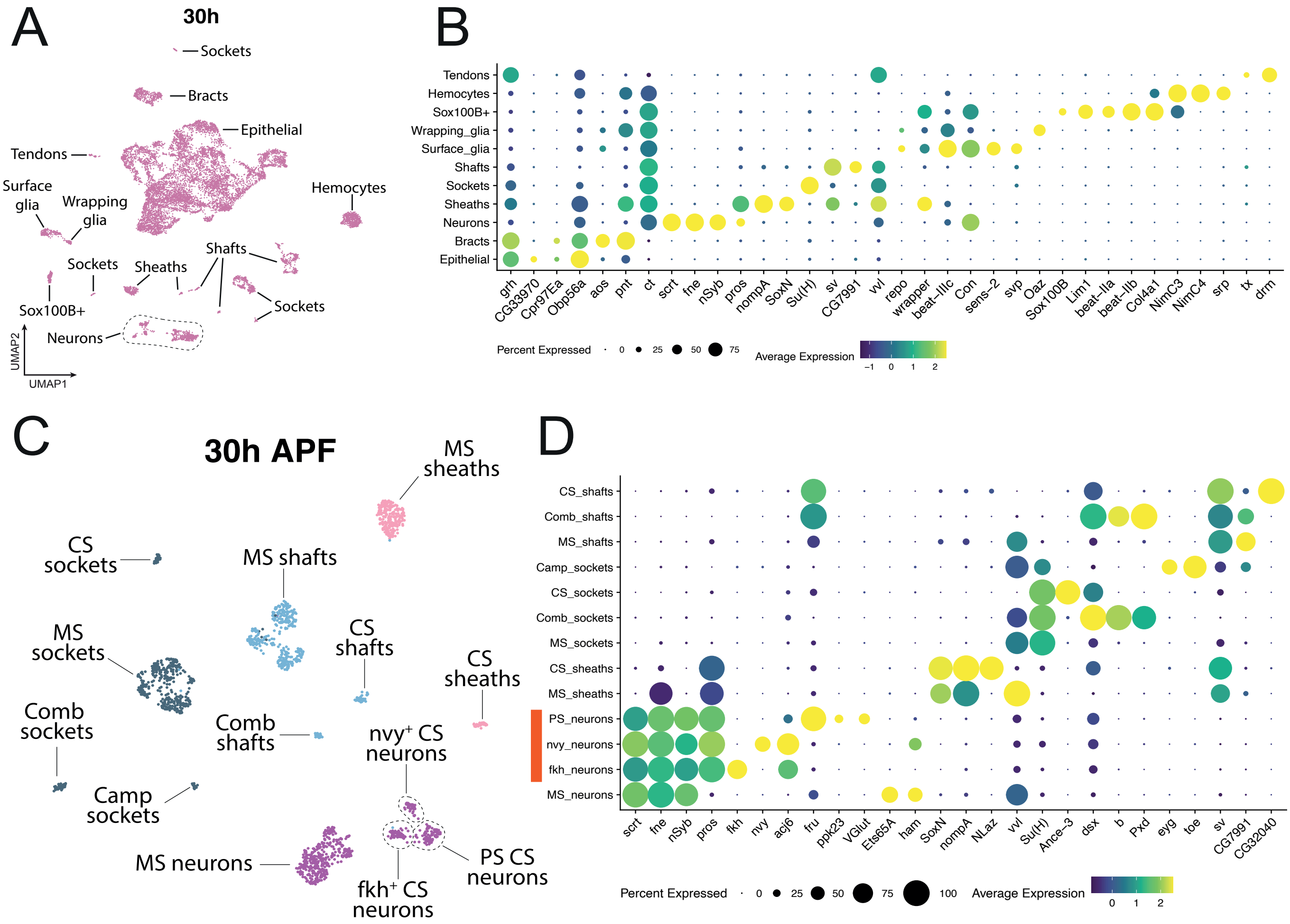


**Supplementary figure 7. Identification of sensory cell populations in the 30h APF male dataset.** (A) A UMAP of the 30h male dataset with major cell populations annotated. Annotation nomenclature follows that used in Hopkins *et al.* (81). (B) Genes enriched for expression in each annotated cluster labelled in (A). (C) The sensory organ lineage cells shown in (A), subsetted out and reclustered with cell types annotated. (D) Genes enriched for expression in each annotated cluster labelled in (C). Note that this dataset was generated from *RAL-517* individuals, so no fluorescent markers are shown. An orange bar denotes subpopulations of chemosensory neuron that were readily identifiable. nvy+ and fkh+ neurons follow the nomenclature used in Hopkins *et al.* (2023). PS neurons are the pheromone-sensing neurons, which at this clustering resolution could not readily be separated into the male- and female-sensing cells seen in Hopkins *et al.* MS=Mechanosensory, CS=Chemosensory, Camp.=Campaniform sensillum.

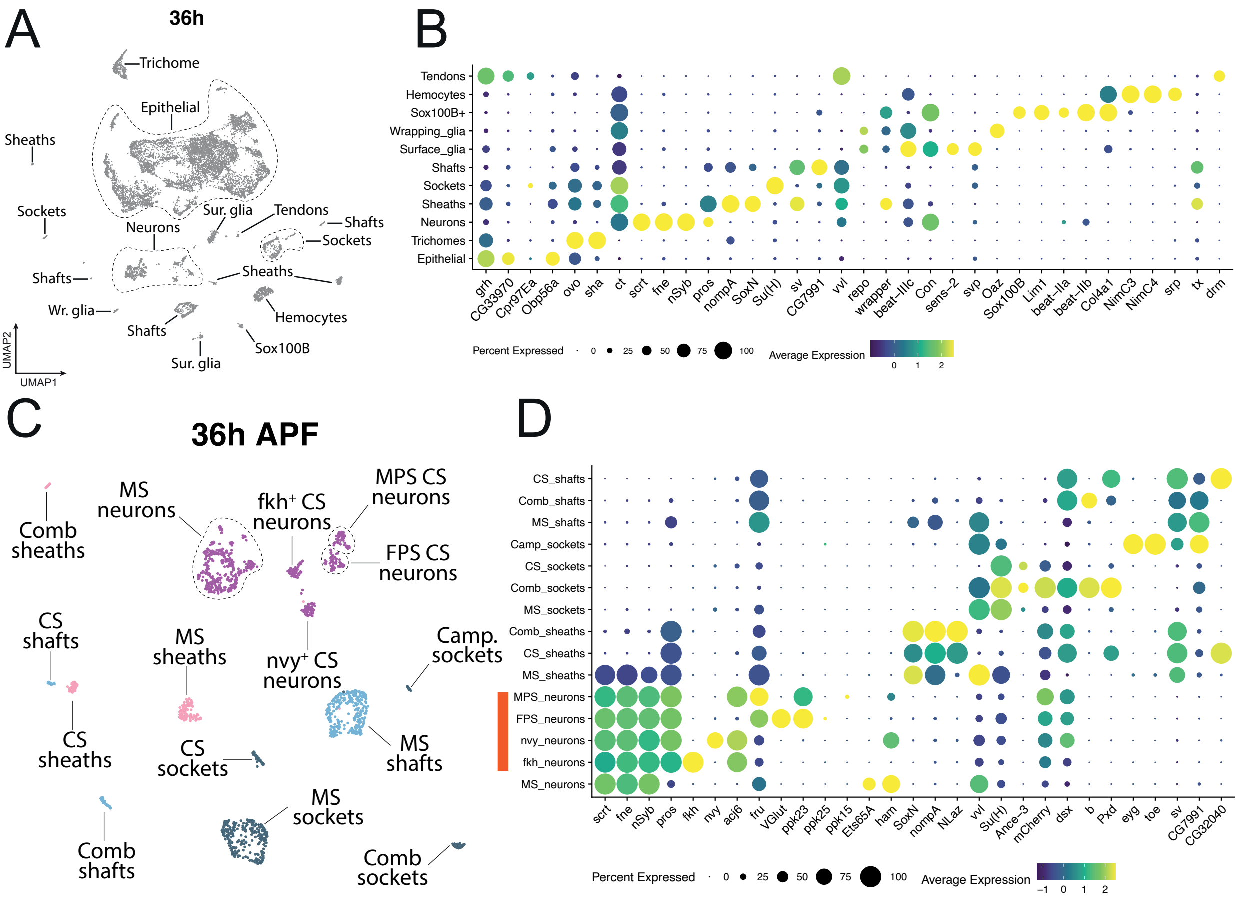


**Supplementary figure 8. Identification of sensory cell populations in the 36h APF male dataset.** (A) A UMAP of the 36h male dataset with major cell populations annotated. Annotation nomenclature follows that used in Hopkins *et al.* (81) with the new addition of trichome cells, which express ovo and sha. Wr. glia = Wrapping glia. Sur. glia = Surface glia. (B) Genes enriched for expression in each annotated cluster labelled in (A). (C) The sensory organ lineage cells shown in (A), subsetted out and reclustered with cell types annotated. (D) Genes enriched for expression in each annotated cluster labelled in (C). Note that this dataset was generated from *dsx*-GAL4 > *UAS-mCherry* individuals, hence the inclusion of *mCherry* in the dotplot. An orange bar denotes subpopulations of chemosensory neuron that were readily identifiable. nvy+, fkh+, male pheromone-sensing (MPS) and female pheromone-sensing (FPS) neurons follow the nomenclature used in Hopkins *et al.* MS=Mechanosensory, CS=Chemosensory, Camp.=Campaniform sensillum.

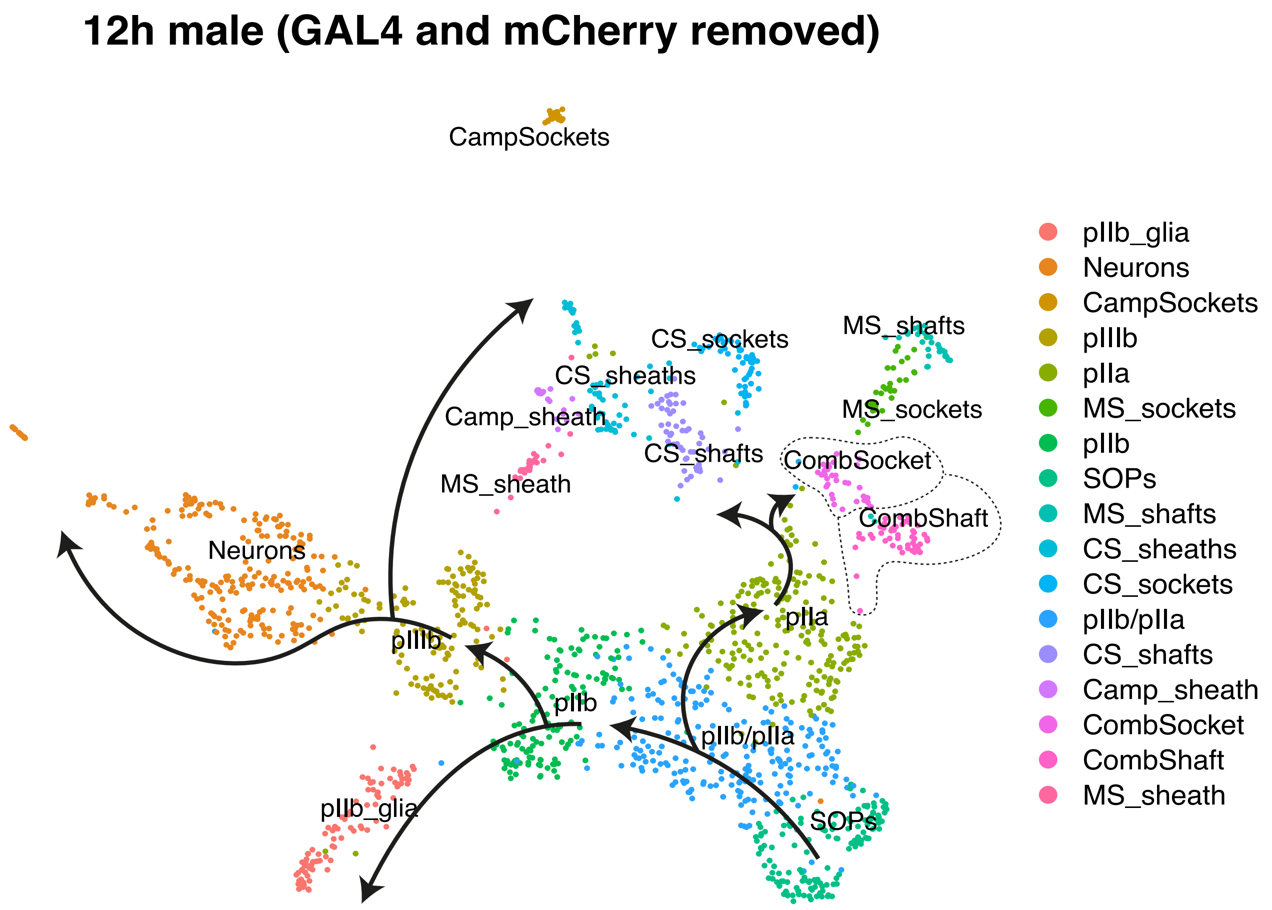


**Supplementary figure 9. The distinct clustering of sex comb cells is not an artifact of the expression of Gal4 and mCherry.** A UMAP of the sensory cells in the 12h APF male dataset after removing *Gal4* and *mCherry* and reclustering. Cells are coloured with the annotation they were given in the dataset that includes *Gal4* and *mCherry*. The comb socket and shaft populations still form distinct clusters indicating that their separation is a product of natural differences in expression. Arrows denote the differentiation trajectories. pIIIb cells divide to form neurons and sheaths, while pIIa cells divide to form sockets and shafts.

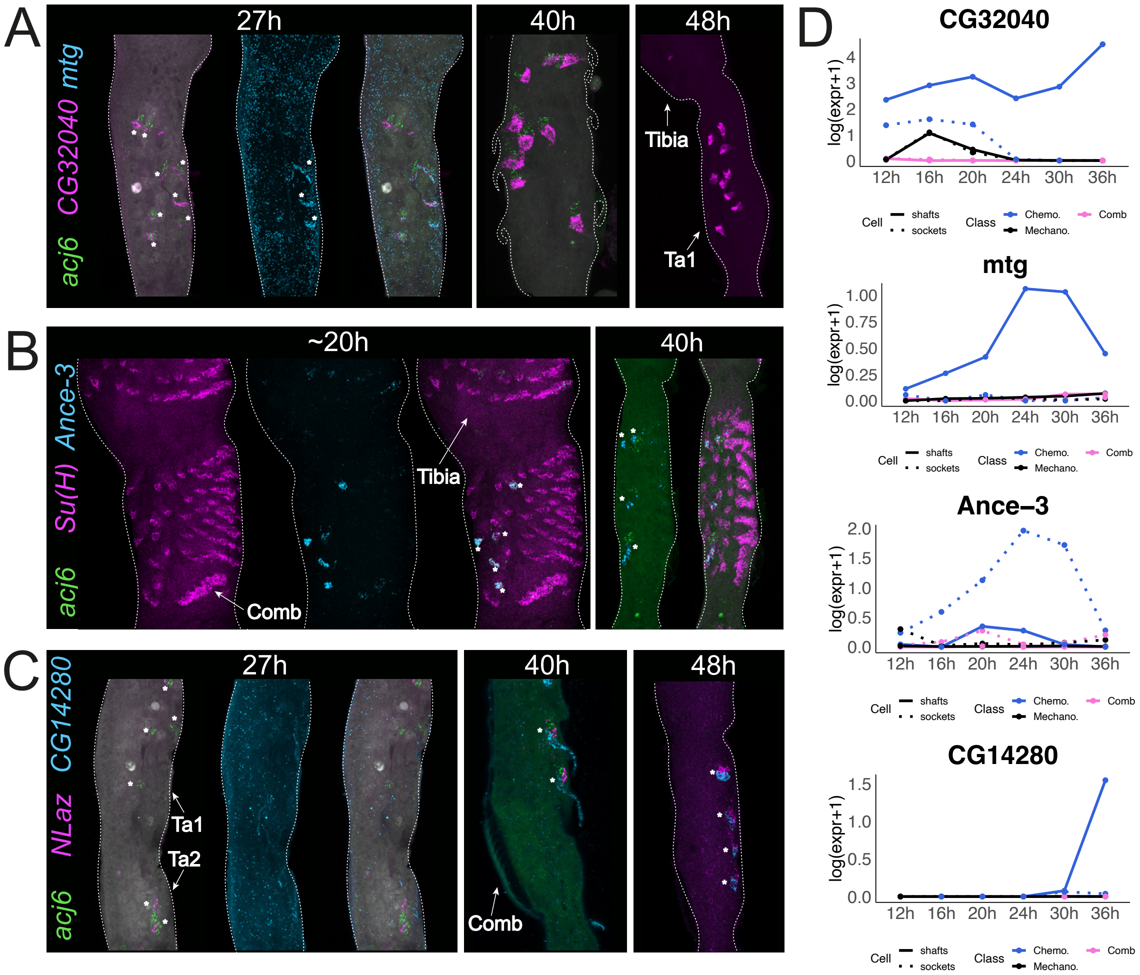


**Supplementary figure 10. Accessory cells in chemosensory bristles express genes that are largely absent from their mechanosensory and sex comb homologues.** (A-C) HCR in situ stainings against genes that are heavily enriched in chemosensory (CS) bristles relative to homologous cells in other bristle classes. *CG32040*, *mtg*, and *CG14280* in shafts, *Ance-3* in sockets, and *NLaz* in sheaths. We use *Su(H)* as a socket marker and *acj6* as a marker of 3 of the 4 neurons that innervate each CS bristle (81). The *mtg* probe shows high background staining throughout the leg but localized staining in the CS shaft is clear. Note that *mtg* staining in (A) is shown only at 27h APF and *acj6* staining in (C) is shown only at 27h APF and 40h APF. Asterisks denote the position of readily identifiable CS bristles in (B) and (C). (D) Time-series expression plots for genes largely specific to CS shaft and socket cells.

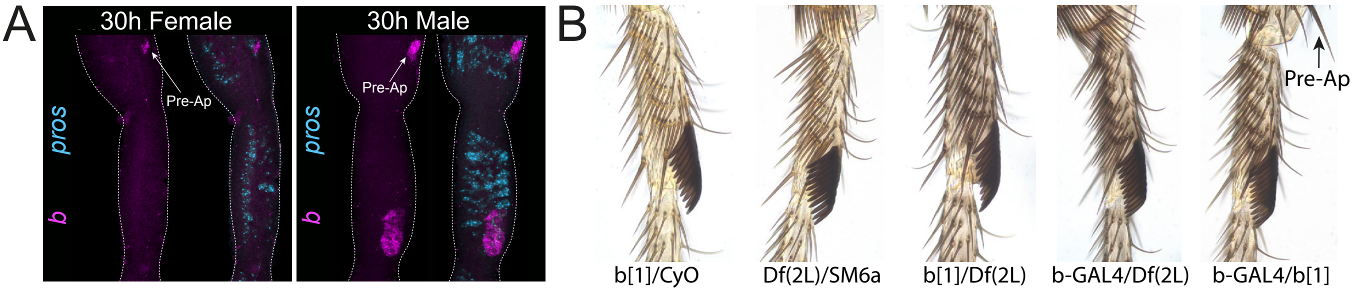


**Supplementary figure 11. *black* is specific to the sex comb in the first tarsal segment but is not required for normal sex comb development.** (A) HCR in situ staining of *b* and *pros* at 30h APF in a male and female first tarsal segment. *pros* is used here as a bristle marker, being expressed in the sheath cell of every bristle and the neurons of chemosensory bristles (81). The position of the sex comb is labeled, as is the likely position of the pre-apical ('pre-ap') bristle in the distal tibia. (B) First tarsal segments from adult males that are transheterozygous for mutant alleles affecting *b*. These alleles are a molecular null (*b[1]*) (82), a hypomorphic GAL4 line (*b-GAL4*)(86), and a deficiency that covers the *b* locus on chromosome arm 2L (*Df(2L)*). Genotypes heterozygous for the molecular null or deficiency line over either the *CyO* or *SM6a* balancer are shown as controls. The pre-apical bristle in the distal tibia is labelled in one image. Note how transheterozygotes show the darker cuticular melanization phenotype known to be associated with *b* nulls (82) but no obvious defects in the sex comb.

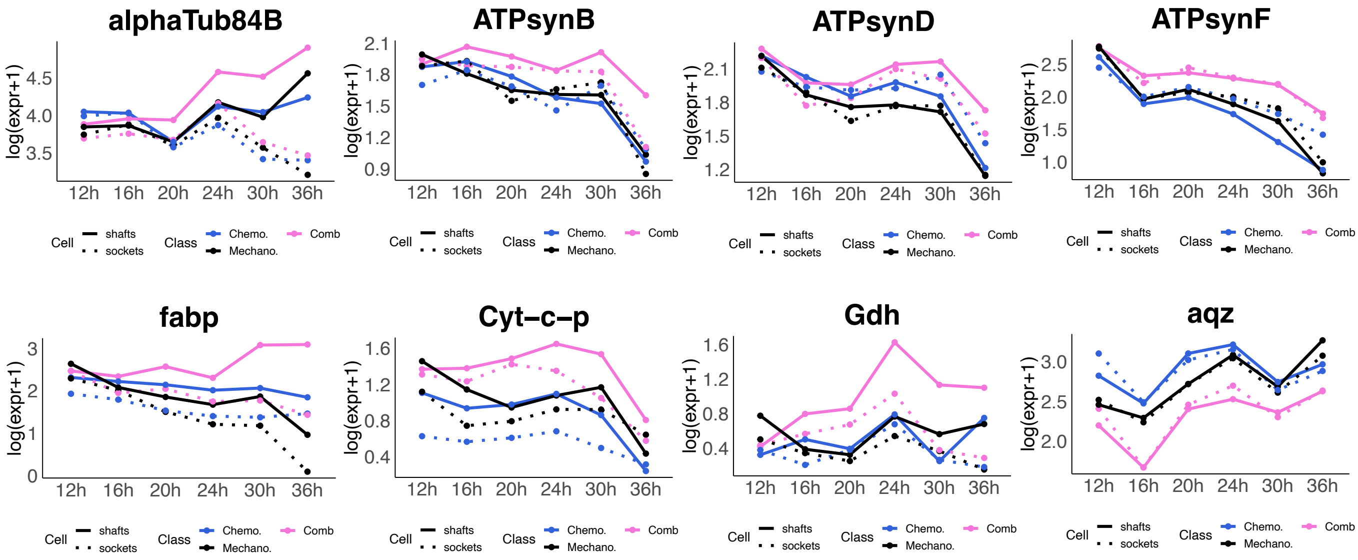


**Supplementary figure 12. Quantitative upregulation of genes in the sex comb and chemosensory bristles.** Time-series expression plots for genes showing significant quantitative expression differences between sex comb and mechanosensory bristle cells, without evidence of heterochrony. The exception is *aqz*, which shows quantitative upregulation in chemosensory rather than sex comb cells.

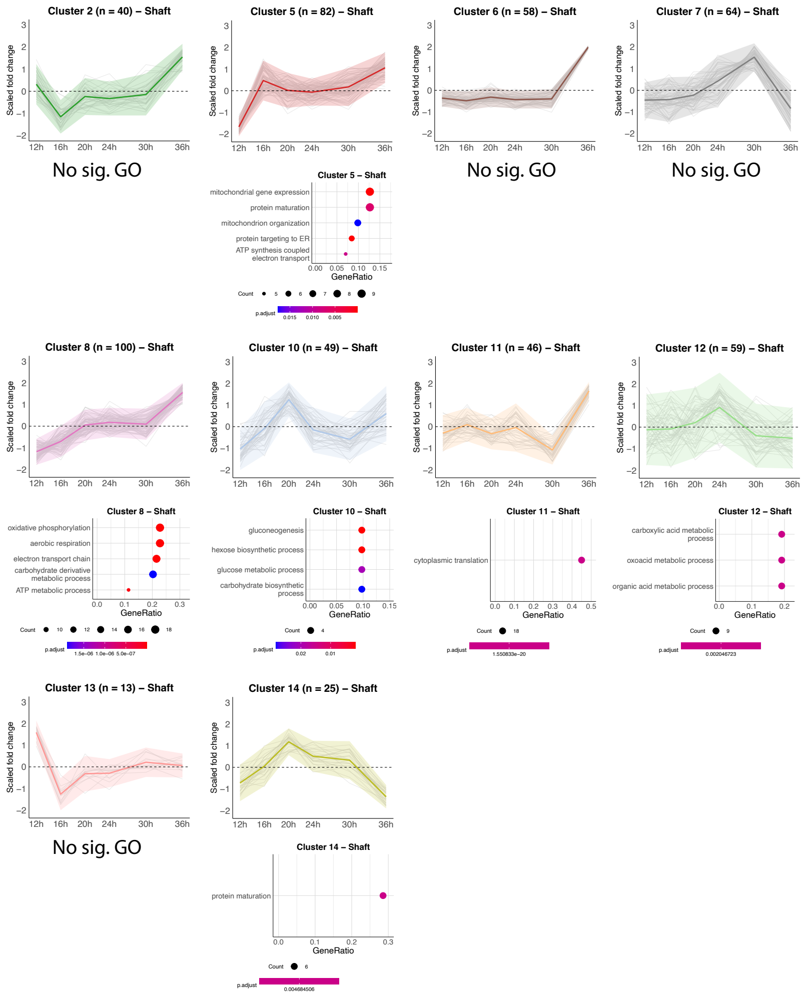


**Supplementary figure 13. Expression trajectories for genes found to be differentially expressed between sex comb shafts and their mechanosensory homologs.** All clusters that are not present in Figure 4 are shown. Genes are grouped based on the cluster they were assigned to in our DPGP analysis. Each expression plot shows the trajectory of each gene in the cluster (gray lines) as well as the cluster mean (thick, colored line). The ribbon denotes ± 2 S.D. The number of genes in the cluster is given in the plot title. The values plotted are the log2 fold changes between sex comb and mechanosensory (MS) cells normalized by Z-score transformation (as in 35). Below each plot is a dot plot showing a selection of up to 5 biological process GO terms found to be significantly enriched in each cluster relative to a background of all the genes detected in MS and comb shaft cells at one timepoint or more.

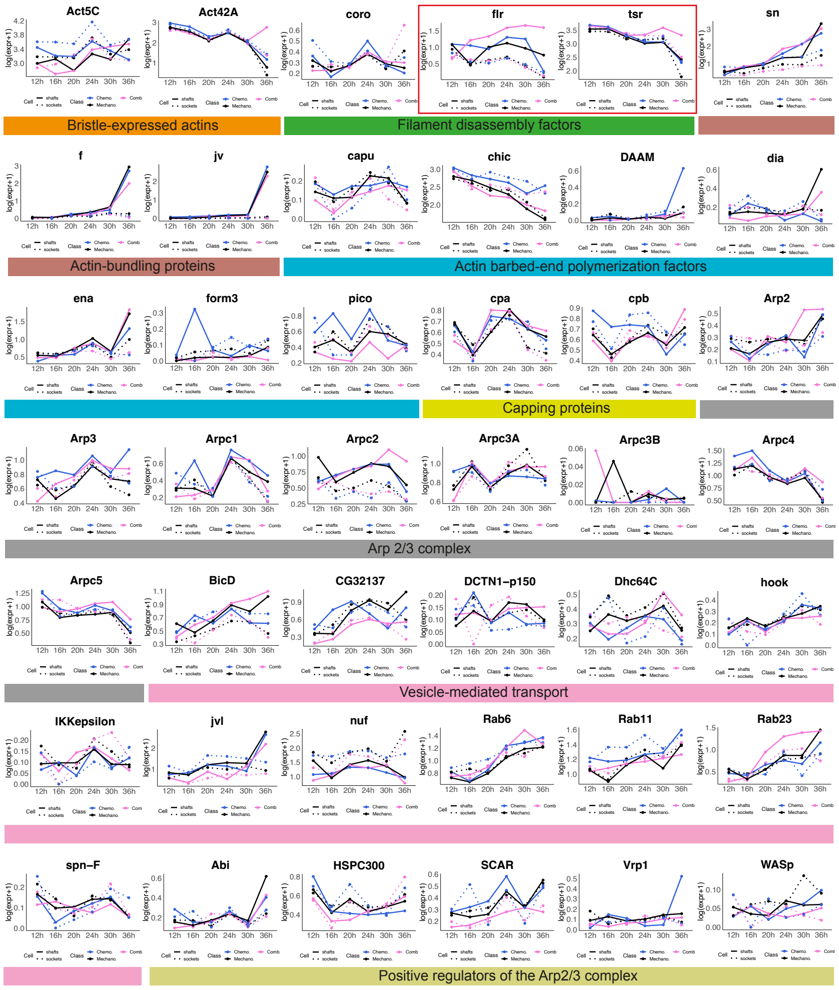


**Supplementary figure 14. Sex comb shafts modulate gene networks that regulate actin filament depolymerization.** Time-series gene expression plots, grouped by the actin-related function each gene performs. Sex comb-enriched expression of the filament disassembly factors *flr* and *tsr* (highlighted with red boxes) provide a potential route through which shaft growth can be accelerated. By pruning and disassembling filaments that are not bundled, these factors ensure the continued availability of actin monomers at the growing tip (38). Note that in many cases, there is no clear difference between shafts and sockets, let alone different classes of shaft.

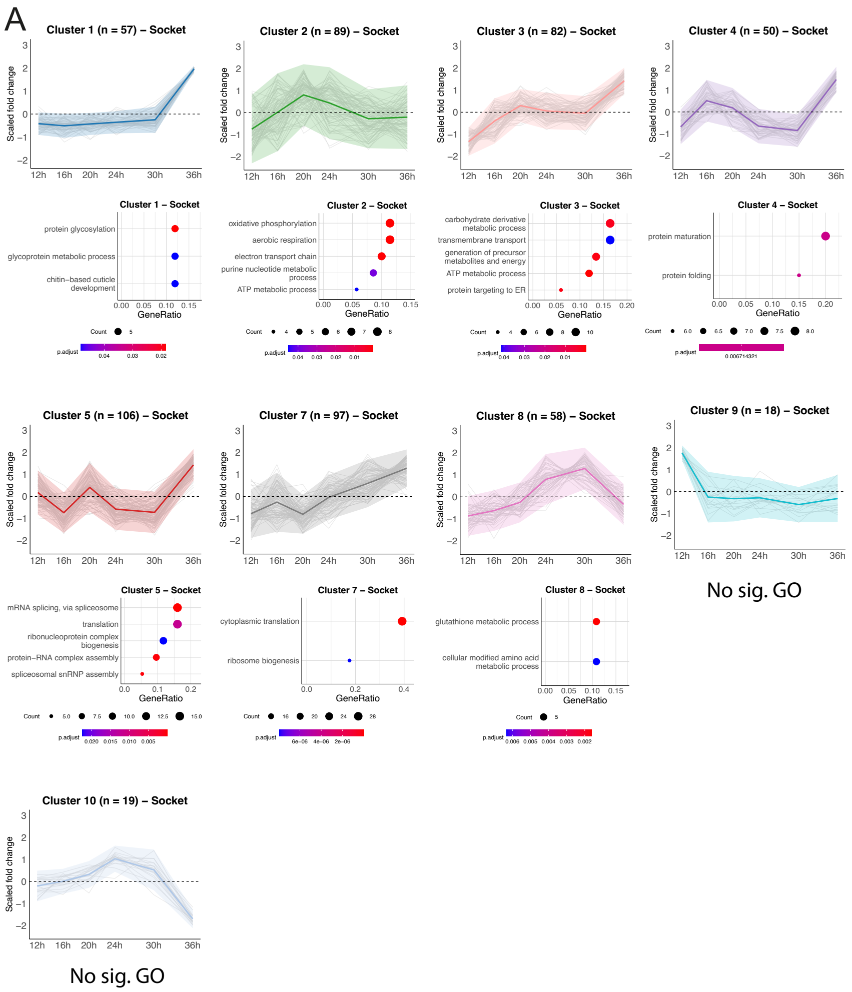


**Supplementary figure 15. Expression trajectories for genes found to be differentially expressed between sex comb sockets and their mechanosensory homologs.** All clusters that are not present in Fig. 5 are shown. Genes are grouped based on the cluster they were assigned to in our DPGP analysis. Each expression plot shows the trajectory of each gene in the cluster (gray lines) as well as the cluster mean (thick, colored line). The ribbon denotes ± 2 S.D. The number of genes in the cluster is given in the plot title. The values plotted are the log2 fold changes between sex comb and mechanosensory (MS) cells normalized by Z-score transformation (as in 35). Below each plot is a dot plot showing a selection of up to 5 biological process terms found to be significantly enriched in each cluster relative to a background of all the genes detected in MS and comb socket cells at one timepoint or more.

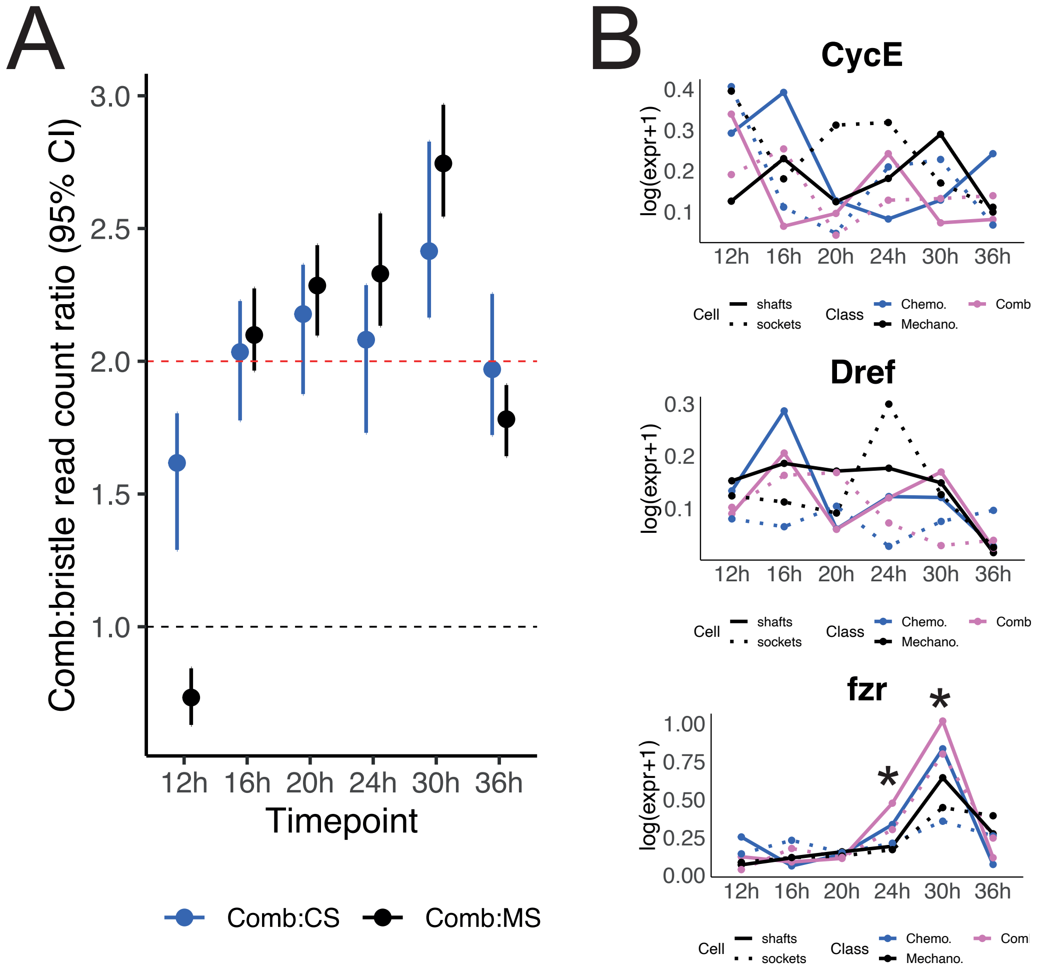


**Supplementary figure 16. Changes in endoreplication dynamics are a key part of the bristle-to-tooth transformation.** (A) The ratio of sex comb shaft to mechanosensory (MS) shaft and chemosensory (CS) shaft read counts at each timepoint. A value of 1 suggests equivalent read counts between, for instance, MS shaft and comb shaft cells, while a value of 2 means that we detect twice as many reads in comb cells. Bootstrapped confidence intervals (2000 iterations) for the ratios are presented. (B) Time-series expression plots for endoreplication-mediating genes we tested for roles in sex comb development using the UAS/GAL4 system. Asterisks denote timepoints where a gene was significantly upregulated in sex comb shafts relative to mechanosensory shafts. Of the three genes we tested—*Dref*, *CycE*, and *fzr*—only *fzr* was significantly upregulated in sex comb cells relative to MS bristles at any time point in our scRNA-seq data, raising the possibility that it might be a direct regulator of the ploidy differences between sex comb and mechanosensory cells. However, we did not see a corresponding increase in MS bristles when overexpressing *fzr* under the control of the bristle driver *Pax2*-GAL4, suggesting that *fzr* alone is insufficient to drive the additional endocycles seen in sex comb cells.

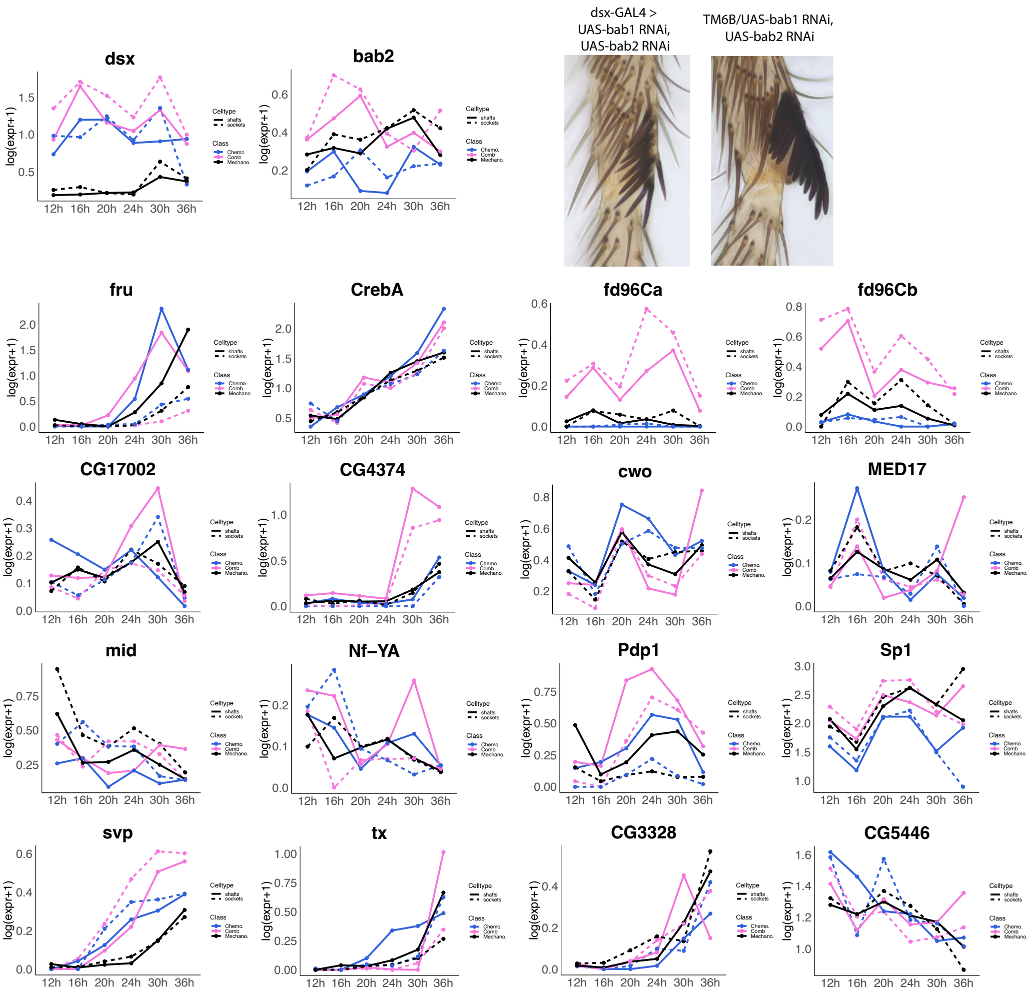


**Supplementary figure 17. Transcription factor knockdowns in the developing sex comb suggest a decentralized genetic pathway.** Expression plots for each transcription factor that we detected as significantly upregulated in sex comb cells at any timepoint. With the exception of *dsx* and the forkhead domain genes *fd96Ca* and *fd96Cb* (which have been tested previously, 51), we used UAS-RNAi constructs to knockdown the expression of each transcription factor in the developing sex comb under the control of *dsx*-GAL4. Only a double knockdown of *bab1* and *bab2* (87) gave a phenotype, showing an incomplete transformation of sex comb teeth towards the thinner, shorter proportions of a mechanosensory bristle.

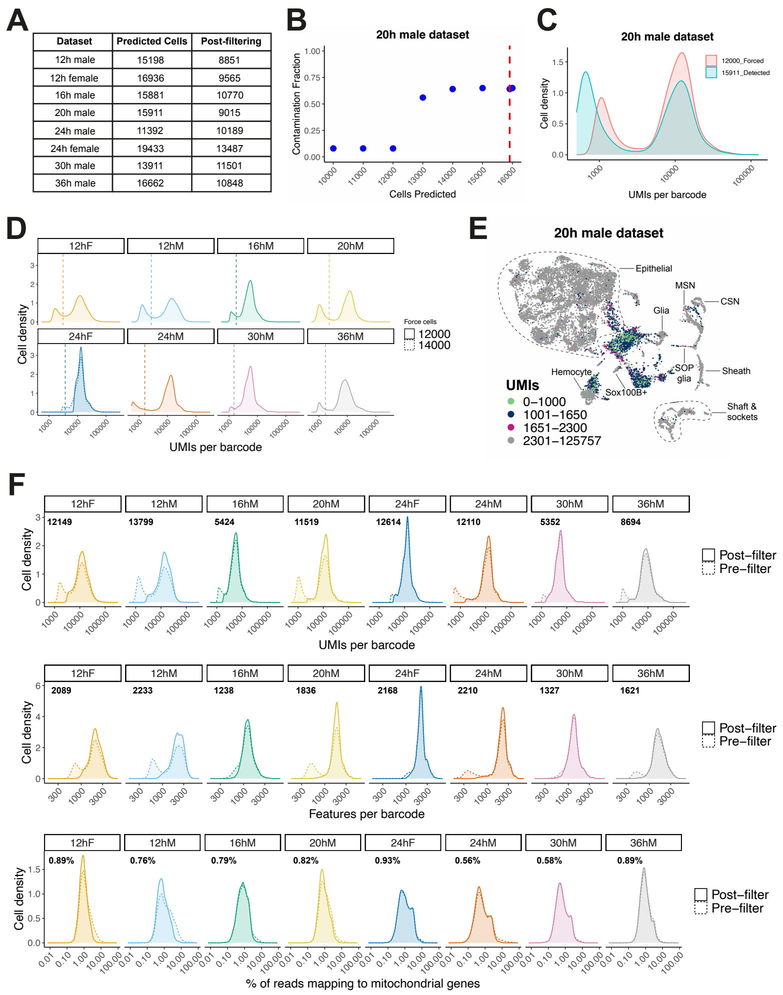


| Dataset | Predicted Cells | Post-filtering |
|---|---|---|
| 12h male | 15198 | 8851 |
| 12h female | 16936 | 9565 |
| 16h male | 15881 | 10770 |
| 20h male | 15911 | 9015 |
| 24h male | 11392 | 10189 |
| 24h female | 19433 | 13487 |
| 30h male | 13911 | 11501 |
| 36h male | 16662 | 10848 |

**Supplementary figure 18. Processing the single-cell data.** (A) The number of predicted cells recovered in each dataset when running CellRanger without ‘force-cells’, alongside the final number of cells in each dataset after filtering cells out based on dataset-specific quality control thresholds. (B) The contamination fraction estimated in the 20h APF dataset when forcing CellRanger to detect set numbers of cells. The dashed red line indicates the number of cells detected (15,911) using default settings (i.e., without force-cells). (C) The distribution of UMIs detected per barcode in the 20h dataset when using default settings or forcing the detection of 12,000 cells. Note the reduction in low barcode ‘cells’ (i.e., putative empty droplets) when instructing CellRanger to force the detection of fewer cells. (D) The distribution of UMIs detected per barcode in each dataset when forcing the detection of 12,000 cells. Note that the female 24h dataset also includes the distribution when forcing the detection at 14,000 cells. We used a higher value for this dataset because cutting at 12,000 lost some of the lower part of the distribution that was retained in all other datasets. (E) A UMAP of the full, force-cells, 20h dataset without any downstream filtering of cells based on quality control metrics. Note how low UMI cells generally form their own cluster or fall between closely associated clusters in UMAP space (e.g., between chemosensory and mechanosensory neuron clusters), suggestive of their aberrant characteristics. (F) The distribution of UMIs, features, and percentage of reads mapping to mitochondrial genes for each of the datasets pre- and post-filtering based on cell-level quality control metrics. The post-filtering median value is provided as a label in each case.
